# Kveik brewing yeasts demonstrate wide flexibility in beer fermentation temperature and flavour metabolite production and exhibit enhanced trehalose accumulation

**DOI:** 10.1101/2021.07.26.453768

**Authors:** Barret Foster, Caroline Tyrawa, Emine Ozsahin, Mark Lubberts, Kristoffer Krogerus, Richard Preiss, George van der Merwe

## Abstract

Traditional Norwegian Farmhouse ale yeasts, also known as kveik, have captured the attention of the brewing community in recent years. Kveik were recently reported as fast fermenting thermo- and ethanol tolerant yeasts with the capacity to produce a variety of interesting flavour metabolites. They are a genetically distinct group of domesticated beer yeasts of admixed origin with one parent from the “Beer 1” clade and the other unknown. While kveik are known to ferment wort efficiently at warmer temperatures, its range of fermentation temperatures and corresponding flavour metabolites produced, remain uncharacterized. In addition, the characteristics responsible for its increased thermotolerance remain largely unknown. Here we demonstrate variation in kveik strains at a wide range of fermentation temperatures and show not all kveik strains are equal in fermentation performance, flavour metabolite production and stress tolerance. Furthermore, we uncovered an increased capacity of kveik strains to accumulate intracellular trehalose, which likely contributes to its increased thermo- and ethanol tolerances. Taken together our results present a clearer picture of the future opportunities presented by Norwegian kveik yeasts and offer further insight into their applications in brewing.

## Introduction

Traditional Norwegian farmhouse ale yeasts, known as kveik, have captured the attention of the brewing community due to their variation from commonly used brewing yeasts (Norland, 1969; Garshol, 2014; Preiss et al., 2018). These poorly characterized yeasts have been used for centuries in western Norway for traditional farmhouse ale brewing characterized by pitching yeast into wort at temperatures in the 28-40°C range and a consequent fast fermentation completing within 1-2 days (Garshol, 2014). The genome sequences of six kveik strains identified them as having an admixed origin, with one parent strain originating from the “Beer 1” clade, as defined by Gallone et al. (2016), and another as of yet unknown parent (Preiss et al., 2018). Genetic and phenotypic characterizations (Preiss et al., 2018) revealed common signs of domestication and promising beer production attributes (reviewed in (Gallone et al., 2018)).

These included efficient flocculation (>80% of strains analyzed) supported by increased copy number variations (CNVs) of various *FLO* genes, efficient consumption of major wort sugars with increased CNVs of maltose and maltotriose metabolic genes (*MAL*), and homozygous loss-of-function single nucleotide polymorphisms (SNPs) in the genes (*PDC1* and *FDC1*) responsible for producing the phenolic off flavour 4-vinylguaiacol, thereby rendering them POF- (Preiss et al., 2018). In addition, kveiks produced a spectrum of fruity esters, notably in the medium chain fatty acid esters, imparting a fruity characteristic to the final product. Finally, phenotypic testing showed remarkable ethanol and thermotolerances for kveik strains, thereby broadening the potential application of these yeasts. It is also clear from Preiss et al. (2018) that these kveik strains, while closely related phylogenetically, are genetically and functionally distinct from each other.

The abovementioned fermentation analyses were performed at 30°C. There is currently no further insight into the range of fermentation temperatures at which diverse kveik strains effectively perform fermentations. The reported ethanol and thermotolerances suggest an increased capacity of kveik strains to combat these stresses during fermentation at higher temperatures, thereby increasing their potential for completing higher-temperature fermentations successfully. Insights into the specific fermentation kinetics and flavour compound production capacities, whether beneficial or undesired, of diverse kveik strains at a wider temperature range are currently lacking.

Ethanol and thermal stress tolerances are polygenic traits that rely on multiple pathways to combat these environmental impacts on the yeast (Caspeta and Nielsen, 2015; Voordeckers et al., 2015; Snoek et al., 2016; Huang et al., 2018). High ethanol concentrations and temperatures disrupt cell wall integrity, membrane fluidity and integrity leading to leaking of cellular content into the environment, increase the toxicity of yeast-produced fatty acids, and denature protein structure to ultimately impact its function. In combination these challenges disrupt the structural integrity of the cell, negatively impacting its metabolism, leading to a decrease in cellular functionality that can lead to cell death. Yeast cells combat increasing ethanol concentrations or high temperatures with intrinsic stress response mechanisms that include stabilizing cell walls and cell membranes, detoxifying its intracellular environment of inhibitory metabolites (e.g., medium chain fatty acids), and increasing the cell’s protein folding capacity (Piper, 1995; Viegas and Sa-Correia, 1997). To this end, molecular chaperones, like the heat shock proteins (Hsp), are induced by environmental stress, including thermal stress, to stabilize protein folding thereby protecting against loss-of-function denaturing (Piper, 1995). Furthermore, the production and accumulation of trehalose, a glucose disaccharide that can function as a reserved carbohydrate produced when glucose becomes limiting, is induced by high temperatures and is known to protect the plasma membrane and proteins from environmental (heat, cold, ethanol) and cellular stresses (ROS/oxidative stress) (Coutinho et al., 1988; Singer and Lindquist, 1998b; da Costa Morato Nery et al., 2008; Eleutherio et al., 2015). The intracellular levels of trehalose are controlled by the coordinated regulation of trehalose biosynthetic and hydrolytic reactions, which are regulated by nutrient signaling and stress response pathways during the life cycle (Singer and Lindquist, 1998b; Eleutherio et al., 2015). In response to stress, trehalose biosynthesis is induced and hydrolysis by trehalases are reduced to accumulate trehalose intracellularly, where it is proposed to bind and stabilize proteins and membranes (Eleutherio et al., 2015; Magalhaes et al., 2018). A high-affinity trehalose transporter, Agt1, exports accumulated trehalose to the periplasmic space, where it is proposed to bind the polar heads of phospholipids in the outer leaflet of the plasma membrane to protect it from environmental stresses (Eleutherio et al., 1995; Eleutherio et al., 2015; Magalhaes et al., 2018). Lastly, yeast cells negate the fermentation inhibitory impacts of MCFAs (e.g., octanoic and decanoic acids), which are amplified by increasing temperature, ethanol concentrations and acidity (Viegas et al., 1989; Viegas and Sa-Correia, 1997), by either expelling them from the cell or converting them to their much less inhibitory ethyl esters (Peddie, 1990; Piper et al., 1998; Cabral et al., 2001; Legras et al., 2010).

To gain further insight into the fermentation characteristics of kveik strains at a range of different temperatures, we compared the fermentation kinetics, sugar metabolisms and flavour compound production profiles of six previously characterized kveik strains (Preiss et al., 2018) to commonly used ale yeasts from Beer 1 and Beer 2 clade yeasts (Gallone et al., 2016). In addition, we investigated the potential mechanism(s) that could contribute to the temperature tolerances of these kveik strains.

## Materials and Methods

### Yeast Strains

We used six previously characterized kveik strains (Preiss et al., 2018) as well as four commonly used commercial ale yeast strains from the Beer 1 and Beer 2 beer yeast families in this study (Supplementary Figure S1A). Cali Ale, Vermont Ale and Kölsch are popular *Saccharomyces cerevisiae* ale strains of the Beer 1 family (Supplementary Figure S1B) (Gallone et al., 2016), which are hypothesized to share a common ancestor with kveik strains (Preiss et al., 2018). St. Lucifer, a Belgian ale yeast of the Beer 2 family (Supplementary Figure S1B), possesses characteristics that are commonly associated with kveik strains, including a high degree of thermotolerance and greater production of fruity flavour compounds but is notably POF+ (Gallone et al., 2016). The kveik strains were previously isolated from batch cultures (Preiss et al., 2018). Phylogenetically Hornindal1 and Laerdal2 are closer to each other and Hornindal2 to Ebbegarden1 (Supplementary Figure S1B). All beer yeast strains were supplied by Escarpment Laboratories (Guelph, Ontario). The commercially available Finnish baking and standard sahti brewing yeast Suomen Hiiva (Katina et al., 2007; Catallo et al., 2020; Magalhães et al., 2021) was used as a control in the trehalose and trehalase experiments.

### Wort Preparation

Wort used for beer fermentations and yeast propagation was obtained from a commercial brewery, Wellington Brewery (Guelph, ON). The hopped wort was prepared using Canadian 2-row malt to an original gravity of 12.5°Plato (1.045 specific gravity). The wort was sterilized prior to use at 121°C for 20 minutes and incubated overnight to the desired fermentation or propagation temperature.

### Propagation and Fermentation

Colonies from YPD plates were inoculated into 100 mL of YPD and grown at 30°C, 170 rpm until cultures reached the pre-diauxic phase of growth. Cell counts were performed using a haemocytometer and cells were transferred into 100 mL of sterilized wort to a targeted cell density of 5 million cells/mL and grown at 30°C, 170 rpm until pre-diauxic phase. These cultures were counted using a haemocytometer and inoculated at a rate of 1.2 × 10^7^ cells/mL into 200 mL of sterilized wort in 250 mL glass bottles fitted with airlocks. These small-scale fermentations were performed in triplicate for 5 days at two colder temperatures (12°C and 15°C), two standard ale fermentation temperatures (22°C and 30°C), and four higher temperatures (35°C, 37°C, 40°C, and 42°C). The bottles were incubated without shaking to best approximate typical beer fermentation conditions. Fermentation profiles were acquired by collecting samples throughout the fermentation process and analyzing changes in specific gravity (SG) using a DMA35v4 portable densitometer (Anton-Paar).

### Beer Composition Analysis

Throughout the fermentation, samples were collected and filtered with 0.45 μm syringe filters prior to metabolite analysis. Flavor metabolite analysis was performed on samples collected at the end of fermentations using HS-SPME-GC-MS (Rodriguez-Bencomo et al., 2012). Samples contained 2 mL of beer, 0.6 g of NaCl, 10 μL of 3-octanol (0.01 mg/mL), and 10 μL of 3,4-dimethylphenol (0.4 mg/mL). 3-octanol and 3,4-dimethylphenol were used as internal standards.

The wort sugar, ethanol and glycerol content were measured using HPLC and a refractive index (RI) detector. The samples collected at the indicated times throughout the fermentation were analyzed using an Aminex HPX-87H column, with 5 mM sulfuric acid as the mobile phase, under the following conditions: flow rate of 0.6 mL/min, 620 psi, and 60°C. Each sample contained 400 μL of filtered sample and 50 μL of 6% (w/v) arabinose as the internal standard.

### Analysis of Trehalose Production and Neutral Trehalase Activity

Colonies were inoculated into 100 mL cultures of YPD and incubated at 30°C with shaking at 180 rpm. Cell samples were collected at the mid-exponential phase of growth and then every two hours until after the diauxic shift to analyze the timing and quantity of trehalose production. Trehalose was extracted from the cells using 0.5 M cold TCA and analyzed via HPLC with an Aminex HPX-87H column at 30°C, 700 psi, and a flow rate of 0.4 mL/min.

Concentration of trehalose was determined relative to a standard curve of known concentrations and expressed as milligrams of trehalose per gram of dry cell weight (DCW). Each sample contained 400 μL of unfiltered sample and 50 μL of 6% (w/v) arabinose as the internal standard.

Culture conditions for trehalase activity assays were identical to those previously described for the trehalose assays. Cell samples were collected two hours post maximum trehalose levels, when trehalase activity is predicted to be highest in all strains. The enzyme kinetics of the hydrolysis of trehalose by neutral trehalase was measured using a stopped assay. Trehalase activity assay was adapted from a previously described assay (Pernambuco et al., 1996). Briefly, cell pellets were collected and resuspended in ice-cold 50 mM MES/KOH (pH 7) + 50 µM CaCl_2_ and crude cell extract was prepared using the glass bead lysis method. Overnight dialysis was performed using a 22 μm Cellu Sep H1 Dialysis Membrane and the dialyzed extract was utilized in stopped trehalase activity assays. The quantity of glucose liberated was determined using a Glucose GOD-PAP Kit (Roche/Hitachi). Trehalase activity was standardized to total protein content using the DC protein quantification technique according to the manufacturer’s recommendations (Bio-Rad).

### Whole-genome sequencing and sequence analyses

DNA was extracted from Kölsch and St. Lucifer as previously described (Preiss et al., 2018). Whole-genome sequencing was performed by the TCAG sequencing facility at Sick Kids (Toronto, Canada). Briefly, an Illumina TruSeq TL paired-end 150 bp library was prepared for each strain and sequenced using an Illumina HiSeq X instrument. This data and WGS data previously generated for the kveik, Cali and Vermont strains (Preiss et al., 2018) were further analyzed. FastQC version 0.11.8 (Andrews, 2010) and fastp version 0.20.1 (Chen et al., 2018) were used to quality-analyze the sequencing reads. Low quality reads and nucleotides were trimmed and filtered using fastp. Filtered reads were aligned against the latest *S*. *cerevisiae* version R64-3-1 genomic reference sequence, which is also known as S288C, using BWA version 0.7.17 (Li and Durbin, 2009). PCR duplicates were marked up and the alignments were sorted and indexed via sambamba version 0.7.1 (Tarasov et al., 2015). Per base depth were calculated via mosdepth version 0.3.0 (Pedersen and Quinlan, 2018) and FreeBayes version 1.2.0 (Garrison and Marth, 2012) was used to perform variant analysis on aligned reads. Variant effect prediction and the gene names were assigned with SnpEff version 5.0e (Cingolani et al., 2012). Copy number variations of genes were estimated based on coverage with Control-FREEC version 11.6 (Boeva et al., 2011). Wilcoxon Rank Sum test (p < 0.05) was performed using the significance script provided by Control-FREEC to evaluate significant copy number variations. Allele frequencies and read depth were plotted to predict the ploidy and aneuploidies for each chromosome.

Phylogenetic analysis was performed using consensus genome sequences generated from variant calls with vcftools (Danecek et al., 2011), which were then aligned to the R64-3-1 genome with minimap2 (Li, 2018). Single nucleotide polymorphisms in the whole-genome alignments were called with paftools.js (included within minimap2), and these were filtered and concatenated using bcftools and bedtools (Quinlan and Hall, 2010). The concatenated SNP matrix was converted into a PHYLIP alignment using vcf2phylip for tree inference with FastTree (Price et al., 2010; Ortiz, 2019). Trees were visualized and annotated in R using the ggtree package (Yu et al., 2017).

### Statistical Analysis

Data were processed and visualized using R. One-way ANOVA with Tukey’s post-hoc test was performed on the HPLC and GC metabolite data using the agricolae package in R (http://www.r-project.org/). The FactoMineR (Lê et al., 2008) and factoextra (http://www.sthda.com/english/rpkgs/factoextra) packages were used to perform Principal Component Analysis.

### Data availability statement

The WGS data of the kveik, Cali Ale and Vermont Ale strains were obtained from NCBI-SRA under BioProject number PRJNA473622 (Preiss et al., 2018) and WGS data for Kölsch and St. Lucifer generated here, are available under the NCBI BioProject accession number PRJNA736724.

## Results

### Kveik displays a wide fermentation temperature range and accelerated fermentation rate

While kveik strains are known to be strong fermenters with increased thermotolerance (Preiss et al. 2018), their fermentation efficiencies at a range of temperatures are poorly characterized. We performed small-scale wort fermentations at eight different temperatures (see Methods) to gain insight into the temperature-dependent fermentation characteristics of six previously characterized kveik strains (Preiss et al., 2018). With a final gravity (FG) ≤ 1.01 as the target for ales (Parker, 2008), the Beer 1 control strains Cali, Vermont and Kölsch, displayed its most efficient attenuation with a SG < 1.01 after only three days at 22°C and 30°C (Figure 1). St. Lucifer (Beer 2) also attenuated best at 22-30°C but never reached an FG < 1.01, regardless of temperature; this is likely due to its inability to properly metabolize maltotriose (Krogerus et al., 2019). These data support moderate fermentation temperatures (22-30°C) as optimal for the control strains. As most traditional kveik fermentations occur at ~30°C or higher (Garshol, 2014), we anticipated their optimal fermentation efficiencies would also be at higher temperatures. 5/6 kveik strains reached a SG < 1.01 within five days at fermentation temperatures between 22-40°C (Figure 1). In most instances, a SG < 1.01 was already achieved within three days at temperatures between 30-40°C, with Hornindal1 (35°C; 37°C), Hornindal2 (35°C; 37°C) and Laerdal2 (30°C) doing so in two days in a temperature-dependent manner.

**Figure 1.**
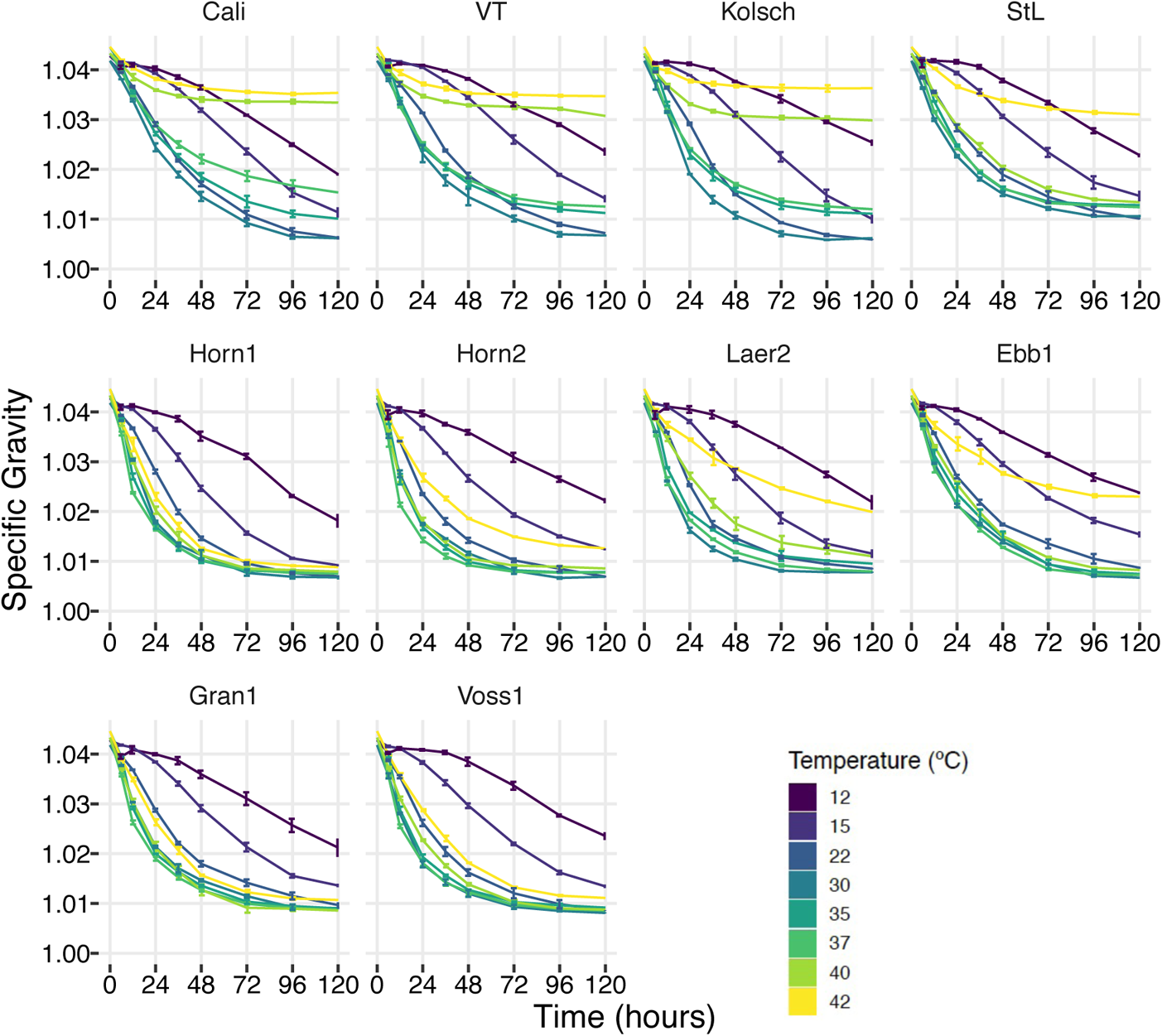
Fermentation profiles at the indicated temperatures between 12°C to 42°C for four commercial *Saccharomyces cerevisiae* ale strains and six Norwegian kveik isolates. Strains were pre-cultured and inoculated into wort as described in the Methods. The fermentation profiles were obtained by recording specific gravities throughout fermentations. Data points represent the mean of biological replicates (n=3) and error bars represent the standard deviation.

These faster attenuations can be partially explained by the rapid initiation of fermentation at these higher temperatures. Within the first 12 hours of fermentation, 5/6 kveik strains already achieved ~30% or more attenuation at 30-40°C, and most notably at 37°C, while the control fermentations were still lagging (Figure 2A). The accelerated attenuation was sustained through 24 and 48 hours (Figures 2B and 2C). Increasing the fermentation temperature to 42°C showed Hornindal1, Granvin1 and Voss1 reaching FG ~ 1.01 or lower, while the Laerdal2 and Ebbegarden1 strains exhibited noticeably impaired fermentations (Figure 1). Thus, kveik strains attenuate with different efficiencies at this higher temperature. By contrast, Beer 1 control strains displayed increasingly weaker attenuation efficiencies as the temperature increased and essentially stopped fermenting after 48 hours at 40°C and 42°C (Figures 1 and 2C). By contrast, St. Lucifer’s attenuation remained consistent at 35°C and 37°C, similar to that of its optimum (Figure 1), with only a slight decrease after 24 and 48 hours (Figure 2B and 2C) and it displayed strong fermentation efficiency at 40°C with an FG similar to that achieved at 35°C (Figures 1 and 2A-C).

**Figure 2.**
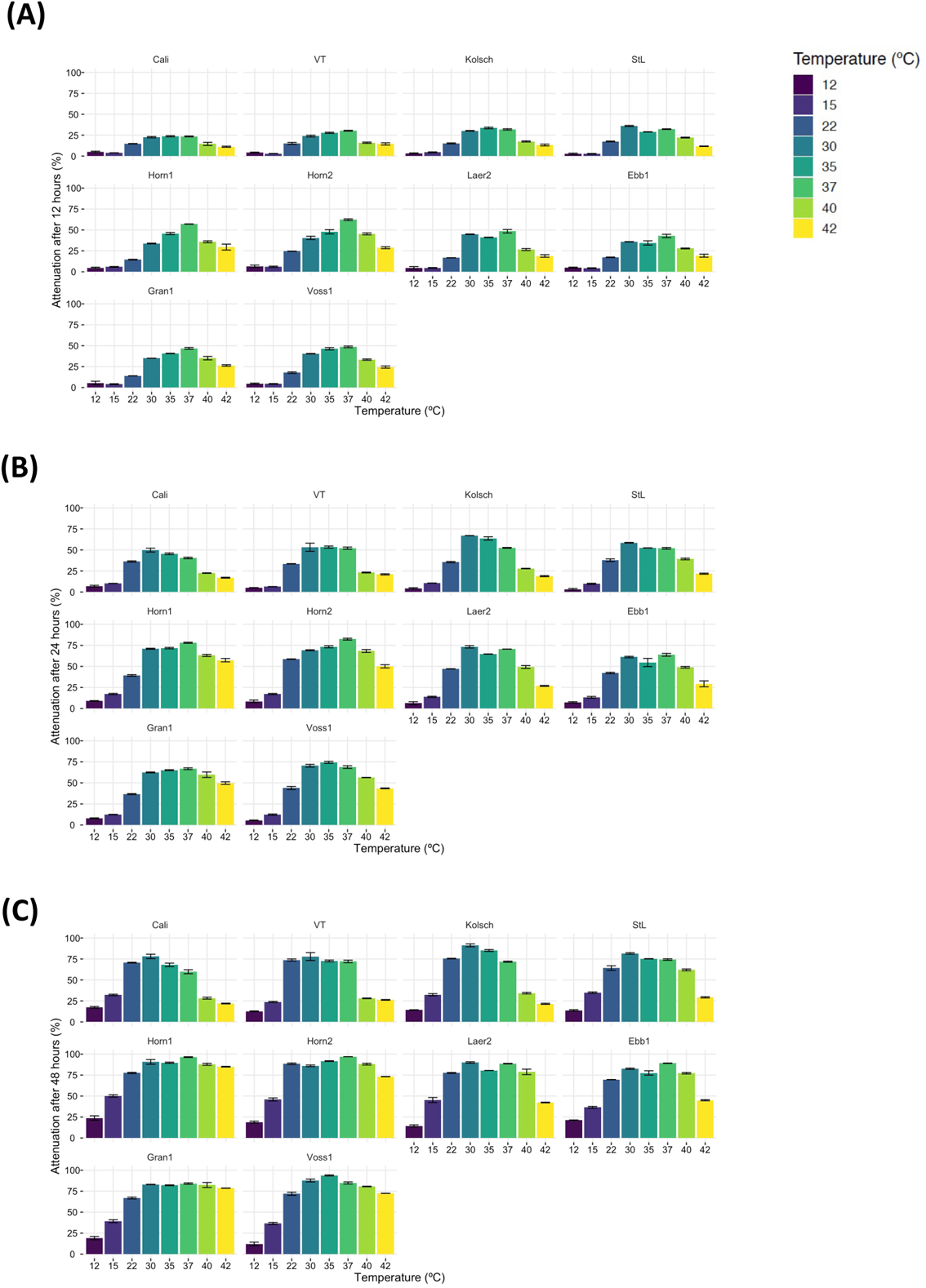
Temperature-dependent changes in specific gravity early in the fermentations presented in Figure 1. Percent attenuation was calculated by determining the difference between the original gravity and the gravity at (A) 12 hours, (B) 24 hours, and (C) 48 hours relative to the final gravity reached by each individual strain at 30°C. Data points represent the mean of biological replicates (n=3), and error bars represent the standard deviation.

In addition to strong fermentation capabilities at higher temperatures, our data also demonstrates that kveik strains possess adequate fermentation efficiencies at lower temperatures. While most kveik strains attenuated similar to the Vermont and St. Lucifer controls at 15°C, Hornindal1 and Laerdal2 completed fermentations similar to the Kölsch and Cali controls, respectively (Figure 1). Again, the kveik strains and in particular Hornindal1, rapidly initiated fermentation at 15°C in comparison to the control strains (Figure 2B and 2C). In addition, Hornindal1 completed a 12°C fermentation within 10 days (Supplementary Figure S3). In combination our data indicates shorter fermentation lag times and faster fermentation rates of selected kveik strains at a wide temperature range between 15-42°C.

### Kveik Exhibit Accelerated Wort Sugar Metabolism

We next focused on wort sugar metabolism to gain further insight into the accelerated attenuation of the kveik strains. Samples collected at the same timepoints of SG measurements (Figure 1) were analyzed by HPLC to monitor wort sugar consumption, and ethanol and glycerol production. The major wort sugar concentrations were maltose (4.7-6 % w/v), maltotriose (1.3-1.5 % w/v), glucose (~1 % w/v), and fructose (~0.2 % w/v) with slight variations depending on the batch of wort. Beer yeasts typically consume the bulk of the available glucose within the first 24h of fermentation, thereby relieving glucose repression that typically inhibits maltose consumption (reviewed in (Stewart and Russell, 1998; He et al., 2014)). Fructose and sucrose are usually present in low concentrations with sucrose being rapidly hydrolyzed in brewing yeasts by invertase even in the presence of glucose, thereby releasing its complement of glucose and fructose for consumption (Meneses et al., 2002; Marques et al., 2016). As fructose is transported and consumed slower than glucose (D’Amore et al., 1989), this often results in a transient spike in fructose early in the fermentation before it is finally consumed within the first three days (Meneses et al., 2002).

We observed differential sugar consumption rates between strains and temperatures. Glucose is rapidly consumed by all strains at their preferred temperature optima with most kveiks consuming >50% glucose within the first 6h progressing to >90% after 12h, while most of the controls were noticeably slower (Figure 3; Supplementary Figure S2). At 42°C, Hornindal1 consumed most of the glucose after 12h, while the control strains and Laerdal displayed the slowest rates (Figure 3; Supplementary Figure S2). In combination these data indicate most of the kveik strains rapidly deplete most of the glucose within the first 12 hours of fermentation at higher temperatures. By contrast, cooler temperatures delayed glucose consumption by all strains (Figure 3; Supplementary Figure S2).

**Figure 3.**
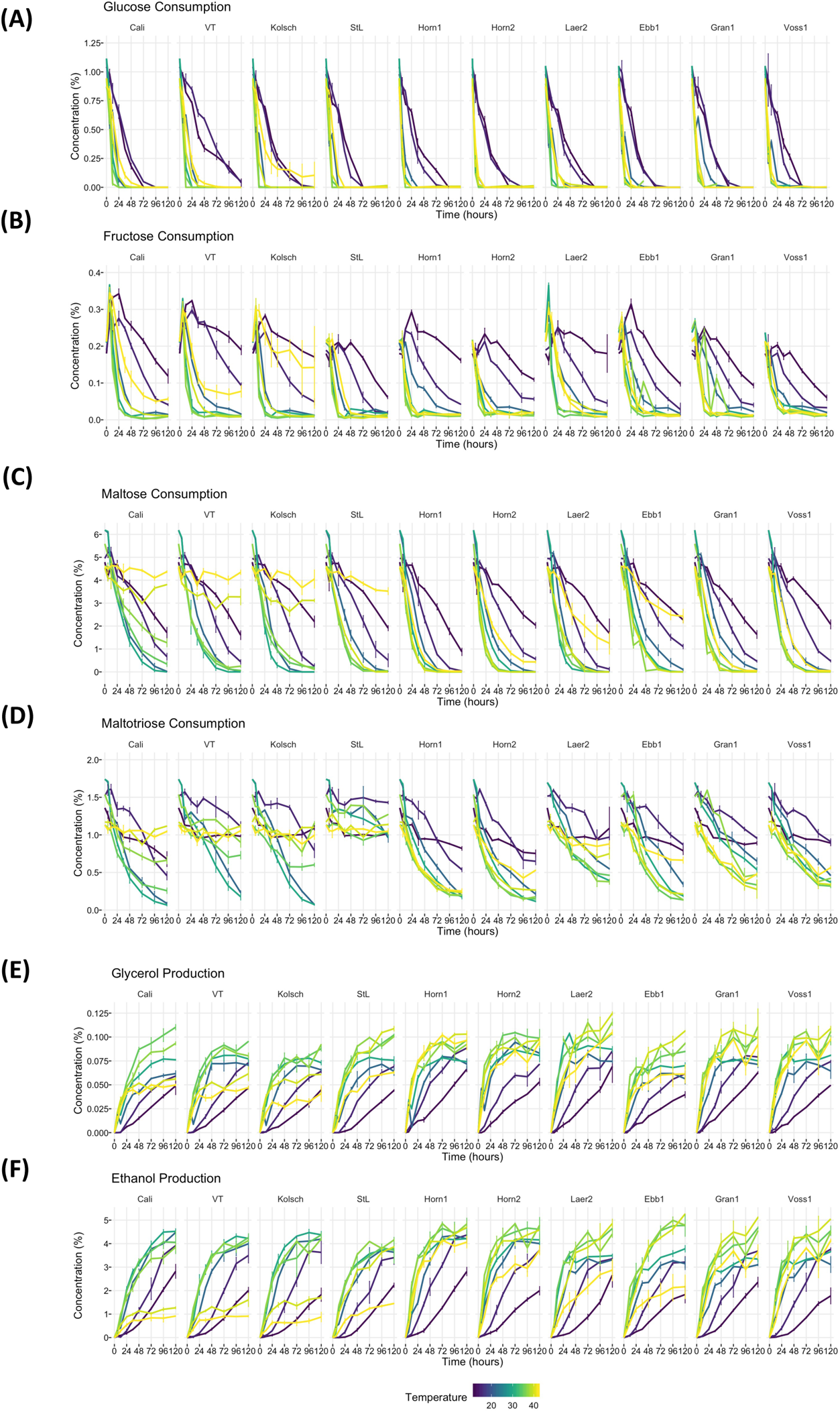
Wort sugar consumption and ethanol and glycerol production of six Norwegian kveik strains and four commercial *Saccharomyces cerevisiae* beer strains during the fermentation in Figure 1. Samples were collected for HPLC analyses at the same timepoints of SG measurements in Figure 1. The concentrations of (A) glucose, (B) fructose, (C) maltose, (D) maltotriose, (E) glycerol, and (F) ethanol were determined by HPLC as described in Methods. Data points represent the mean of biological replicates (n=3) and error bars represent the SD.

All the strains metabolized fructose to negligible residual levels by the end of the fermentation at 22-40°C, while the kveik and St. Lucifer strains also did so at 42°C (Supplementary Figure S2). By contrast, most strains showed slow fructose consumption at colder temperatures (120h at 12°C) (Figure 3; Supplementary Figure S2). However, prolonged fermentation at 12°C showed all strains, with the exception of Hornindal2 (86.5%) consumed >90% of the fructose after 10 days (Supplementary Figure S3). These data indicate that the kveik strains consume fructose to negligible amounts at a broad temperature range.

The overall fermentation rates of maltose and maltotriose are known to correlate with its respective transport efficiencies (Kodama et al., 1995; Meneses et al., 2002; Rautio and Londesborough, 2003; Alves et al., 2007). Also, maltose transport and hence fermentation rate in ale yeasts is much more efficient at moderate (20°C) than cooler temperatures (Rautio and Londesborough, 2003). As maltose is the main sugar in wort, we expected the faster fermentation rates of kveik would be explained by faster maltose consumption. Maltose consumption was initiated within the first 6 hours by all the strains at almost all temperatures. As expected, the kveik strains depleted maltose noticeably faster than the controls with most kveiks consuming at least 30% of this sugar after 12 hours and 75-90% after 24 hours of fermentation in the 30-37°C range (Figure 3; Supplementary Figure S2). Maltose uptake usually starts when approximately 60% of the available glucose has been consumed (D’Amore et al., 1989). The kveik strains depleted glucose faster than the control strains during the first 6 hours of fermentation (Supplementary Figure S2), thereby potentially relieving glucose repression on maltose metabolism faster than the controls. Generally, the controls and kveik strains consumed maltose faster at their respective fermentation optima; the controls were most efficient at 30°C while the kveik yeasts consumed the bulk of the available maltose within 24 hours at 30-37°C (Supplementary Figure S2). Previous WGS analyses of these kveiks revealed considerable amplifications of various *MAL* loci of which the *MAL3x* locus (*MAL31* permease, *MAL32* maltase and *MAL33* transcription factor) is estimated to be present 11 or more times depending on the strain (Preiss et al., 2018). These CNVs in kveik may therefore contribute to its increased maltose consumption; however, much variability in the rates of maltose consumption exist among the kveik yeasts (Supplementary Figure S2; Maltose 12h and 24h). Finally, the Beer 1 strains were deficient in maltose consumption at 40-42°C, while St. Lucifer could deplete maltose at 40°C, but not 42°C. The kveik yeasts, however, depleted maltose at these higher temperatures with only Ebbegarden1 and Laerdal2 showing deficient maltose consumption at 42°C.

All the strains tested showed slower maltose consumption at lower temperatures (12-15°C), likely because glucose consumption was also slower at these lower temperatures (Supplementary Figure S2) and maltose transport is known to be slower at cooler temperatures (Rautio and Londesborough, 2003). Interestingly, most of the kveik yeasts showed effective maltose consumption, with some strain variation, after 120 hours at 15°C (Supplementary Figure S2). Fermentations at 12°C with a prolonged incubation (10 days) revealed kveiks can deplete maltose at these cooler temperatures (Supplementary Figure S3). In combination, these data confirm the rapid consumption of maltose by kveik strains at their optimal fermentation temperatures and the ability of some, but not all, strains to effectively consume maltose at higher (42°C) and lower (15°C) temperatures. The rapid and efficient consumption of these wort sugars are also reflected in ethanol produced during the fermentation (Supplementary Figure S4; Ethanol). These findings expand the potential applications of kveik in brewing and other fermentation industries.

The rate limiting step in both maltose and maltotriose consumption is its transport into the cell (Kodama et al., 1995; Meneses et al., 2002; Rautio and Londesborough, 2003; Alves et al., 2007). α-glucosides, including maltose and maltotriose, are actively transported across the plasma membrane by H^+^-symporters (Kodama et al., 1995; Rautio and Londesborough, 2003; Stambuk et al., 2006). Unlike maltose, maltotriose does not have its own high-affinity transporter(s); instead, some maltose transporters can also transport maltotriose, but with much lower affinity. Maltotriose transport and its consequent consumption is therefore usually delayed to the latter stages of wort fermentations (Han et al., 1995; Day et al., 2002; Salema-Oom et al., 2005). Nonetheless, increasing the fermentation temperature (15 to 21°C) is known to increase the rate of maltotriose utilization in beer yeasts (Zheng et al., 1994). While kveik strains have been shown to consume maltotriose at 30°C (Preiss et al., 2018), its efficiency of consuming this sugar at a range of temperatures is not known. Here, maltotriose was consumed slower and less completely than the other wort sugars by all strains and fermentation temperature had a pronounced impact. While none of the kveik strains depleted maltotriose completely after 120 hours of fermentation, Hornindal1 and Hornindal2 initiated maltotriose usage the fastest and at the widest temperature range (30-42°C) (Figure 3, Supplementary Figure S2). Hornindal1 (80-90%), Hornindal2 (73-93%), and Ebbegarden1 (80-90%) consumed more maltotriose (30-40°C) than Granvin1 (68-76%) and Voss1 (71-78%) after 120 hours (Supplementary Figure S2), while Laerdal2 showed inefficient maltotriose consumption at all temperatures. Finally, the kveik strains, except Laerdal2, consumed ~50% of the maltotriose at 42°C with Hornindal1 being able to use 78% of the sugar (Supplementary Figure S2). At the colder temperatures, maltotriose consumption was slower. Again, the kveik strains showed some variation in maltotriose consumption at 12°C after a 10-day fermentation; Hornindal1 consumed 84% of the maltotriose while Laerdal2 and Granvin1 were least effective with ~55% consumption (Supplementary Figure S3). Thus, while the kveik strains can achieve a FG ~1.01 after 10 days in a 12°C fermentation, most, with the exception of Hornindal1, will leave a significant amount of maltotriose at the end of the fermentation.

The main transporter responsible for maltotriose uptake in beer yeast is *AGT1* (Alves et al., 2007; Alves et al., 2008). From the WGS data of the kveik strains (Preiss et al., 2018) we identified two heterozygous frameshift mutations in *AGT1*; the 1175_1176insT is present in Ebbegarden1 and Laerdal2, while 1772delA (His591fs) is present in Voss1 and Hornindal2 (Supplementary Table S2). Also, two gained stop codons (heterozygous) were identified among the kveik strains; 491AAG>TTA (Leu164*) is again present in Laerdal2 and Ebbegarden1, while 1236C>G (Tyr412*) is heterozygous in Hornindal2 (Supplementary Table S2). These SNPs suggest heterozygous loss of function of Agt1 in 4/6 kveik strains.

Beer 1 control strains initiated maltotriose consumption effectively at their optimal fermentation temperatures (22-30°C) and proceeded to deplete most of maltotriose after 120 hours of fermentation (Figure 3; Supplementary Figure S2). By contrast, these control strains showed inefficient maltotriose consumption (<75%) outside their fermentation temperature optima, with almost no consumption beyond 37°C (Supplementary Figures S2 and S3). By contrast, St. Lucifer used ~25% of the available maltotriose at 22-30°C within the first 12 hours and then stalled consumption at all temperatures (Figure 3, Supplementary Figure S2). This specific strain carries a frameshift mutation after Glu395 (Glu395fs) in *AGT1*, impacting its ability to efficiently metabolize maltotriose (Gallone et al., 2016). In addition, St. Lucifer also has a deletion in the *STA1* promoter, which reduces production of the Sta1 glucoamylase (Krogerus et al., 2019) thereby enabling only slow maltotriose utilization.

Similar to ethanol, the kveik strains produced glycerol faster than the controls through the first 24 hours of the fermentation, with clear differences in glycerol concentrations present in the 35-40°C temperature range (Supplementary Figure S4; Ethanol and Glycerol, 12 hours and 24 hours). Glycerol production is induced by moderate heat shock and it serves as a mechanism to maintain the intracellular redox balance to regenerate the NAD^+^ consumed during anaerobic fermentation (Omori et al., 1996; Arroyo-Lopez et al., 2010; Du et al., 2012). The increased sugar metabolism and higher preferred fermentation temperature could be linked to the initial faster production of glycerol by the kveik strains. After 120 hours all the strains tested displayed similar lower levels of glycerol at the 12-30°C fermentation range, while the control strains, with the exception of St. Lucifer, produced lower levels of glycerol at temperatures beyond 37°C. By contrast, the kveik strains, with the exception of Ebbegarden1, produced higher levels of glycerol at 35-42°C after 120 hours (Supplementary Figure S4; Glycerol, 120 hours). In combination, these data suggest the higher fermentative capacities of the kveik strains at the higher temperatures could in part be attributed to the slightly higher levels of glycerol produced at these temperatures.

### Kveik flavour profiles adjust in a temperature-dependent manner

In addition to fermentation efficiency and flexibility, the flavour metabolite production of brewing yeast is a highly relevant outcome for brewers. We next determined if the volatile flavour profiles developed by the kveik strains differed from each other and the control strains in a manner dependent on the fermentation temperature. We performed HS-SPME-GC-MS analyses on the samples collected at the end of each fermentation described in Figure 1. We measured metabolites that can be separated into four major groups: fatty acids, ethyl esters, alcohols, and acetate esters (Supplementary Table S1).

PCA biplot analysis of the GC-MS metabolite data revealed clustering amongst the strains in a temperature-dependent manner (Figure 4). Most of the kveik strains, with the exception of Ebbegarden1, tend to group together while the controls Cali, Kölsch, and Vermont cluster together. St. Lucifer tends to cluster separately from these Beer 1 controls and was among the kveik strains (Figure 4). In addition to the clustering seen based on strain type, temperature also played a significant role in how the strains clustered. For example, the flavour metabolites produced by kveiks clustered independently based on temperatures. Higher temperatures of 35-42°C clustered the kveik strains in metabolite profiles distinct from those produced at 15-22°C (Figure 4). The Beer 1 control strains were largely distinct from the kveiks at most temperatures tested. These data suggest distinct temperature-specific flavour compound production profiles for kveik yeast strains that are different from those produced by the control strains, with the exception of St. Lucifer.

**Figure 4.**
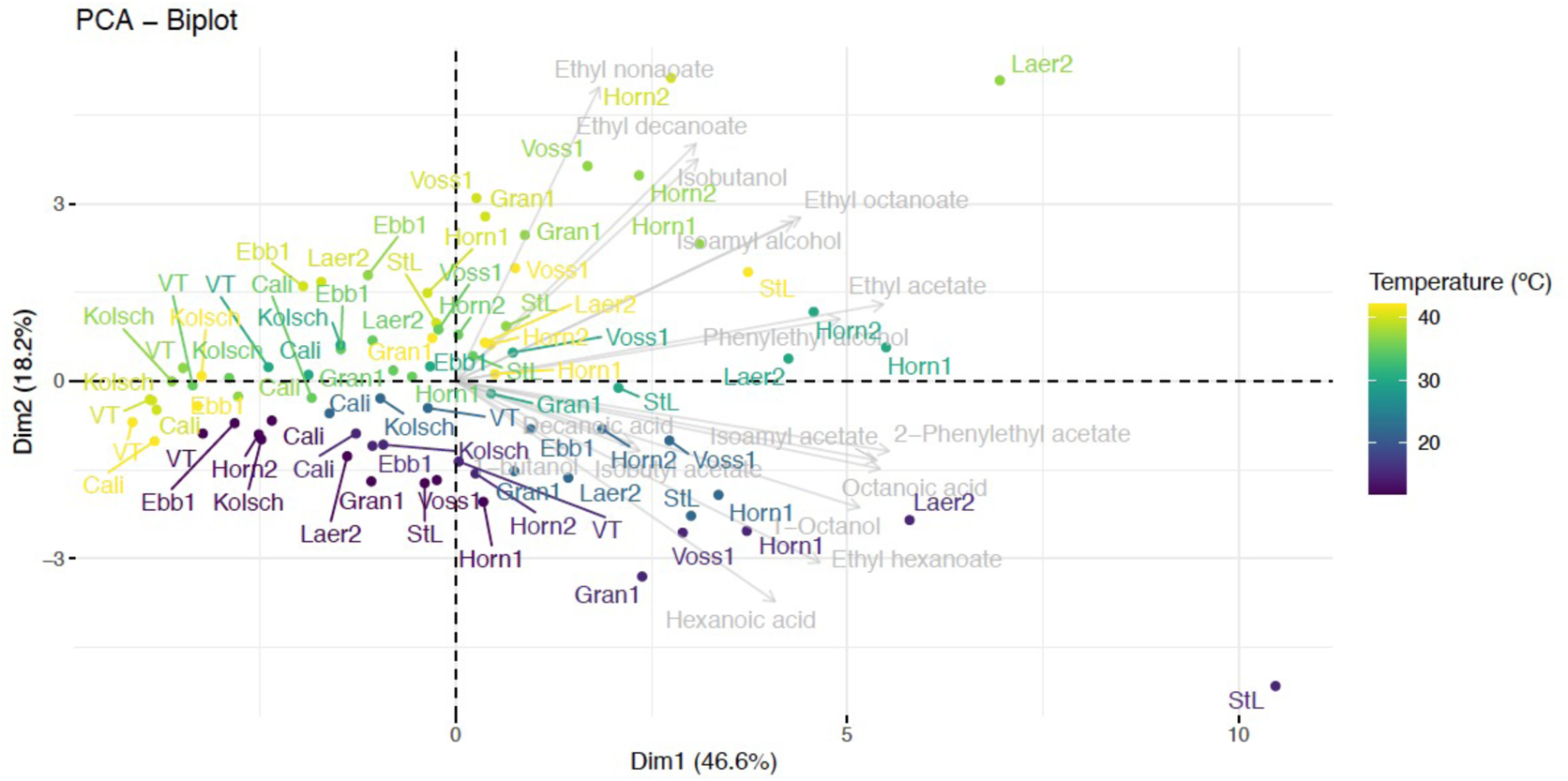
PCA analysis of volatile flavour compounds produced at different fermentation temperatures. Samples collected at the end of the fermentations in Figure 1 were analyzed by HS-SPME-GC-MS. PCA was performed on individuals and variables using the R package FactoMineR. Any missing values were replaced by the column mean. Data points represent the mean of biological replicates (n=3) and are labelled with the strain name. Specific fermentation temperatures are colour-coded.

The kveik strains, with the clear exception of Ebbegarden1, showed enhanced flavour compound production related to medium chain fatty acid (MCFA) (C_6_ – C_10_) metabolism across most of the fermentation temperatures compared to the Beer 1 control strains (Supplementary Figure S5, Supplementary Table S1). Octanoic acid is the most abundantly produced MCFA at each temperature, with concentrations noticeably higher than those of hexanoic and decanoic acid (the latter often not detected), for all strains (Supplementary Figure S5). Octanoic acid is produced at above-threshold concentrations (3 ppm; (Engan, 1972; Meilgaard, 1982; Verstrepen et al., 2003; Comuzzo et al., 2006)) at all the fermentation temperatures by most of the kveik strains and St. Lucifer. The data shows some kveik strains, like Hornindal1 and Laerdal 1, produce octanoic acid at higher concentrations while others, like Ebbegarden1, do not. Interestingly, octanoic acid was present at the highest concentrations in fermentations performed at <30°C, while its abundance clearly decreased at warmer temperatures. While hexanoic acid is not produced above its threshold (8 ppm), the kveik strains and St. Lucifer produce more of this fatty acid than the Beer 1 controls and notably so at <30°C (Supplementary Figure S5, Supplementary Table S1).

Yeast condense MCFAs (acyl-CoA) and ethanol as precursors into ethyl esters (Ayrapaa and Lindstrom, 1977; Taylor and Kirsop, 1977; Dufour et al., 2003). Increasing fermentation temperatures are known to increase the production and release of ethyl esters (Suomalainen, 1981; Saerens et al., 2008a). Saerens et al. (2008) showed ethyl octanoate and ethyl decanoate concentrations increased as the temperature increased in the 20-26°C range while ethyl hexanoate remained constant. Three of the ethyl esters we measured, ethyl hexanoate (pineapple, tropical; threshold: 0.21 ppm), ethyl octanoate (tropical, apple, cognac; threshold: 0.9 ppm), and ethyl decanoate (apple; threshold: 0.2 ppm) (Engan, 1972; Meilgaard, 1982; Verstrepen et al., 2003; Comuzzo et al., 2006) were consistently produced at above-threshold concentrations by the kveiks and St. Lucifer and generally in higher amounts compared to Cali, Vermont, and Kölsch (Supplementary Figure S5, Supplementary Table S1). This was particularly striking for ethyl octanoate at temperatures >30°C where Laerdal2, Voss1, and Hornindal1 produced the highest amounts in a strain and temperature-specific manner (Supplementary Figure S5, Supplementary Table S1). Conversely, Ebbegarden1 produced the least amount of these esters, likely due to the lower amounts of precursors produced. In addition, the kveiks generally produced similar or slightly higher levels of ethyl octanoate than the Beer 1 controls at 12-22°C. By contrast, all strains increased their ethyl decanoate production as the fermentation temperature increased; noticeably for the kveik strains and St. Lucifer (Supplementary Figure S5, Supplementary Table S1). Ethyl decanoate was lowest at 12°C and peaked at 30-42°C range for Laerdal2, Voss1, Hornindal1 and St. Lucifer depending on the specific strain. In combination these findings suggest the optimum fermentation temperatures for ethyl octanoate and ethyl decanoate production are in the 30-40°C range. In combination these findings suggest strain-specific fermentation temperature optima to achieve targeted MCFA-related flavour compound production.

MCFAs are produced by the fatty acid synthase (FAS) complex in a manner dependent on acetyl-CoA carboxylase (*ACC1*) (Ayrapaa and Lindstrom, 1977; Taylor and Kirsop, 1977; Wakil et al., 1983; Dufour et al., 2003; Marchesini and Poirier, 2003). *FAS1* and *FAS2* encode the beta and alpha subunits of the FAS complex, respectively, and both these genes are known to participate in MCFA and ethyl ester synthesis (Furukawa et al., 2003). In addition, a point mutation in *FAS2* has been shown to enhance octanoic acid synthesis (Nagai et al., 2016). Also, the long-chain fatty acyl CoA synthetase Faa1 is proposed to detoxify the inhibitory effects of MCFAs (Legras et al., 2010). We identified several SNPs (homozygous or heterozygous) in *AAC1*, *FAS1*, *FAS2* and *FAA1* that are unique to either the kveik or the respective control strains (Supplementary Table S2), suggesting differential impacts of these mutations on the functionality of the respective enzymes. For example, seven SNPs in *ACC1* are unique in 4/6 or more kveik strains. Some of these, like 5510A>C (Asn1837Thr) and 2995G>A (Val999Ile) are homozygous in some and present in high copy number in other kveik strains (Supplementary Table S2). *FAS1* contain six SNPs present in 3/6 or more kveik strains. Some of these, like Ala620Ser and Glu1216Asp, are homozygous in 3/6 kveik strains. Interestingly, the heterozygous Lys879Met SNP in 3/6 kveik strains destroys a known ubiquitination site in Fas1 (Swaney et al., 2013) that could impact its intracellular abundance. *FAS2* carries three SNPs unique to 3/6 or more kveik strains. While these SNPs are mostly heterozygous, the Met802Val mutation is homozygous in Voss1. Interestingly, the SNPs we identified in *FAA1* were present only in the closely related Hornindal2 and Ebbegarden1 (Supplementary Table S2; Supplementary Figure S1).

Two acyl-CoA:ethanol *O*-acyltransferases, Eeb1 and Eht1, are responsible for ethyl ester synthesis with Eeb1 being the major contributor (Saerens et al., 2006; Saerens et al., 2008a). In addition, phospholipase B with acyltransferase activity (Plb2) was shown to be a strong effector of ethyl octanoate production (Merkel et al., 1999; Steyer et al., 2012). SNP analyses of *EEB1*, *EHT1*, and *PLB2* identified several heterozygous variants that are unique to either the kveik or the control strains (Supplementary Table S2), suggesting different capacities to produce ethyl esters. Interestingly, some SNPs (e.g., *EEB1* 793G>A, Gly265Arg) were unique to closely related Hornindal2 and Ebbegarden1, while other SNPs absent from these two strains were uniquely present in the other kveik strains tested (e.g., *EEB1* 742G>A, Val248Ile). Lastly, yet other SNPs were uniquely homozygous in Hornindal2 and Ebbegarden1, while carrying low frequencies of this mutation in other kveiks (e.g., *EHT1* 355G>A, Asp119Tyr; *PLB2* 1828A>G, Ile610Val), suggesting genetic variations that could impact ethyl production among kveik strains. Whether either of these SNPs impact ethyl ester production is currently not known.

All strains tested produced higher alcohols, like butanol (0.5 ppm), isoamyl alcohol (70 ppm), isobutanol (100 ppm), and phenethyl alcohol (100 ppm) (Engan, 1972; Meilgaard, 1982; Verstrepen et al., 2003; Comuzzo et al., 2006), at levels below their respective sensory thresholds (Supplementary Table S1). Nonetheless, some kveik strains (with the exception of Ebbegarden1) and St. Lucifer clearly produced phenylethyl alcohol at higher concentrations than the Beer 1 control strains; this was also the case for isoamyl alcohol at higher temperatures (Supplementary Figure S5, Supplementary Table S1). Interestingly, we identified 1-octanol, described as having a pleasant citrus-like aroma (Hunter and Moshonas, 1966), was present at levels at or above the sensory threshold only at 15°C (Cali and Vermont) and 22°C (Cali), while the kveik and St. Lucifer strains produced it above the sensory threshold at a wider temperature in a strain-dependent manner (Supplementary Table S1).

We also measured four acetate esters and found isoamyl acetate (banana; 1.2 ppm), and to a lesser extent phenylethyl acetate (rose, honey; 3.8 ppm) (Engan, 1972; Meilgaard, 1982; Verstrepen et al., 2003; Comuzzo et al., 2006) were produced above the sensory threshold (Supplementary Figure S5, Supplementary Table S1). The Beer 1 controls produced isoamyl acetate above threshold at its fermentation optima (15-30°C), but failed to do so for phenethyl acetate. By contrast, the kveik strains and St. Lucifer produced isoamyl acetate above threshold at a wider temperature range including fermentations performed >30°C. Phenethyl acetate, on the other hand, was only produced above threshold by Hornindal1, Hornindal2, Laerdal2 and St. Lucifer without any defined temperature pattern. The abundances of these acetate esters again fluctuated in a strain and temperature-dependent manner.

Phenylethyl acetate is produced through the shikimate pathway that utilizes *ARO1*,*2*,*3*,*4*,*7*,*10* and *PHA2* to produce phenylacetyl aldehyde which is reduced to phenylethyl alcohol by alcohol dehydrogenases (*ADH1-6*, *SFA1*) (Larroy et al., 2002; Dickinson et al., 2003; Holt et al., 2019). Alternatively, the transamination of phenylalanine by the aromatic aminotransaminases I/II (Aro7/Aro8) also yields phenylethyl alcohol, which is finally converted to phenylethyl acetate by the alcohol acetyltransferases Atf1 and Atf2 (reviewed in (Holt et al., 2019)). These latter two enzymes, the branched-chain amino acid aminotransferase Bat2, the Pdc1, 5, 6 and Aro10 decarboxylases and the *ADH* genes are key components in isoamyl acetate production (reviewed in (Holt et al., 2019)). In addition, isoamyl acetate-hydrolyzing esterase Iah1 impacts isoamyl acetate levels (Fukuda et al., 1998). We uncovered SNPs unique to the kveik strains in *ARO1*, *ARO2*, *ARO4*, *ARO10*, *PHA2*, *ADH1*, and *ADH6* (Supplementary Table S2). Some kveik strains carried homozygous mutations in these genes (*ARO1* 3286A>C, Ile1096Leu in Ebbegarden1; *ARO4* 42G>T, Lys14Asn in Laerdal2 and Hornindal1; *ARO10* 551C>T, Ala184Val in Hornindal1). In addition, 4/6 kveik strains carry a heterozygous loss-of-function mutation in *ADH6* (235delC, Lys80fs) and 6/6 kveiks strains share mutations in *ADH1* and *ADH4* that are present in St. Lucifer, but absent from Beer 1 strains (Supplementary Table S2). Furthermore, unique loss-of-function mutations in *ATF1* (1577A>G, 526STOP gained; homozygous in Laerdal2 and heterozygous in Voss1) and *ATF2* (1275delG, Val427fs; heterozygous in Ebbegarden1 and Hornindal1) were previously reported along with the observations that these mutations did not impact acetate ester production at 30°C (Preiss et al., 2018). Importantly, these mutations predict diminished rather than absent alcohol acyltransferase activity. Additional SNPs unique to the kveik strains and absent from the controls are present in *ATF1*, *ATF2*, *BAT2* and *IAH1* (Supplementary Table S2). The notably unique SNPs present in all six kveik strains include *ATF1* 642A>C (Lys214Asn; homozygous in Hornindal2 and Laerdal2) and *BAT2* 748A>G (Ser250Gly; homozygous in Hornindal2, Granvin1 and Laerdal2). Interestingly, while *BAT2* 865A>T (Thr289Ser) is heterozygous in the Kölsch strain, it is homozygous in 5/6 kveik strains. Whether or not these specific alleles impact acetate ester production in kveik, is currently unknown.

### Kveik displayed increased cell viability at higher temperatures

The kveik strains were clearly metabolically more active than the control strains at higher temperatures, while the Beer 1 controls showed significant deficiencies (Figure 3; Supplementary Figure S2). As higher temperatures impact yeast cell viability (Lindquist, 1986; Davidson et al., 1996), we measured the survival of these yeasts at 35-42°C. The Beer 1 control strains were impacted at all the temperatures tested as they showed decreased cell viabilities of <75% at 37°C and above, but variable susceptibilities to higher temperatures; e.g., Kölsch showed 50% viability at 40°C, while Cali were <5% viable (Figure 5). St. Lucifer was much more temperature tolerant, only displaying significant cell death at 42°C. The increased cell mortalities of the controls correlate well with their observed decreases in fermentation efficiency, sugar consumption and ethanol production at higher temperatures (Figures 1–3). By contrast, all the kveik strains, except for Laerdal2, showed no noticeable cell death at 35-37°C (Figure 5) and these viabilities were reflected in the increased fermentation efficiencies and sugar metabolisms at these temperatures (Figures 1–3). Interestingly, as the temperature increased further, the Hornindal1, Granvin1 and Voss1 strains showed more cell survival (>90% at 40°C and >75% at 42°C) than the remaining kveik strains (Figure 5). Hornindal2 still had ~75% viability at 40°C, but this decreased substantially to only 25% at 42°C and impacted its attenuation efficiency (Figure 1), and maltose and maltotriose consumptions at this temperature (Figure 3). Laerdal2 and Ebbegarden1 were clearly more sensitive to 40-42°C (Figure 5), that similarly had a detrimental impact on their attenuation, and maltose and maltotriose consumption capacities (Figures 1 and 3). Interestingly, chromosomal copy number analyses revealed the three most tolerant kveik strains had 4n+1 aneuploidies of Chr IX (Supplementary Table S3). In combination, our data revealed differential tolerances among kveik strains to high temperatures which correlated with attenuation and sugar consumption efficiencies.

**Figure 5.**
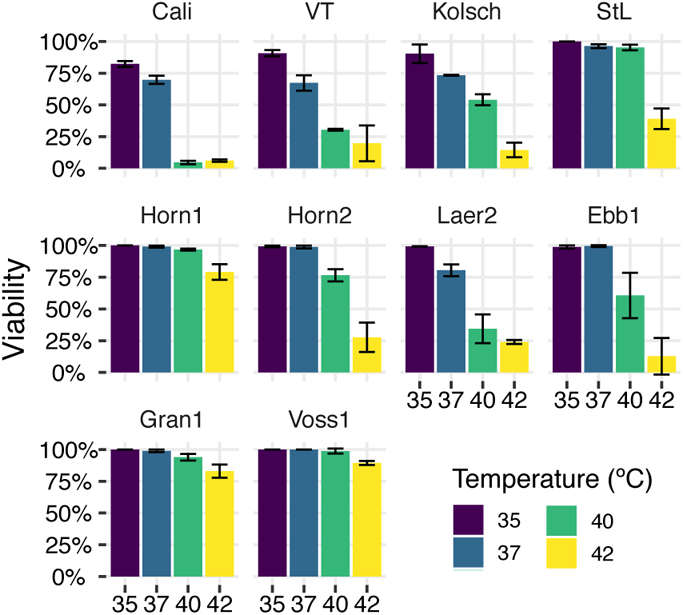
Cell viability following heat treatment. Single colonies of the indicated four commercial ale yeast controls and six Norwegian kveik strains were inoculated into YPD and cultured for 24 hours with shaking at 170 rpm at 35°C, 37°C, 40°C and 42°C. Cell viability was ascertained via staining with Trypan Blue. Error bars represent SD, n=3.

### Kveik Exhibit Enhanced Trehalose Accumulation and Reduced Trehalase Activities

Trehalose is used as a storage carbohydrate and a stress protectant in yeast (Eleutherio et al., 2015). It is typically not produced during early exponential growth on glucose; however, it starts to accumulate in the diauxic phase when glucose becomes depleted as cells transition into stationary phase (Lillie and Pringle, 1980; Francois and Parrou, 2001). Importantly, trehalose is also rapidly produced, even in early exponential phase, in response to stress, and its accumulation is pivotal to combat cold, heat and ethanol stress (Hottiger et al., 1987; Eleutherio et al., 1993; De Virgilio et al., 1994; Hottiger et al., 1994; Bandara et al., 2009). To gain further insight into the tolerance of the kveik strains to the higher fermentation temperatures, we measured trehalose production in selected kveik and control yeasts, and the baker’s yeast Suomen Hiiva as a trehalose-producing control. Yeasts were grown in YPD at optimal growth (30°C) and increased (37°C) temperatures, respectively, and cell samples were collected up to 72 hours to monitor the timing and magnitude of trehalose synthesis and accumulation. Trehalose was undetectable during the early and mid-exponential phases of growth at 30°C. When cells approached the diauxic shift and most available glucose in the media was depleted, trehalose synthesis was induced as seen by the rapid increase in intracellular trehalose during diauxic growth (Figure 6A). Strikingly, the initial trehalose peaks were substantially higher for the kveiks than the control strains ranging from ~100 mg trehalose/g DCW for Hornindal2 and Granvin1, to ~125 mg trehalose/g DCW for Hornindal1 and Voss1 (Figure 6A). By contrast, the control strains produced noticeably less trehalose. Not only did the kveik strains produce more trehalose, they also did so much faster than the control beer strains. The kveik strains all produced >90 mg trehalose/g DCW after 22 hours of growth, before the control strains produced any detectable trehalose (Figure 6A). Additionally, Suomen Hiiva, which as a bread and standard sahti brewing yeast should accumulate intracellular trehalose (Catallo et al., 2020; Magalhães et al., 2021), peaked at 63 mg trehalose/g DCW. The kveik strains therefore produced >1.5-2-fold trehalose and in a shorter timeframe than the beer control yeasts as an initial response at 30°C. This discrepancy in trehalose production suggests misregulation of or modifications to trehalose metabolism in the kveik strains.

**Figure 6.**
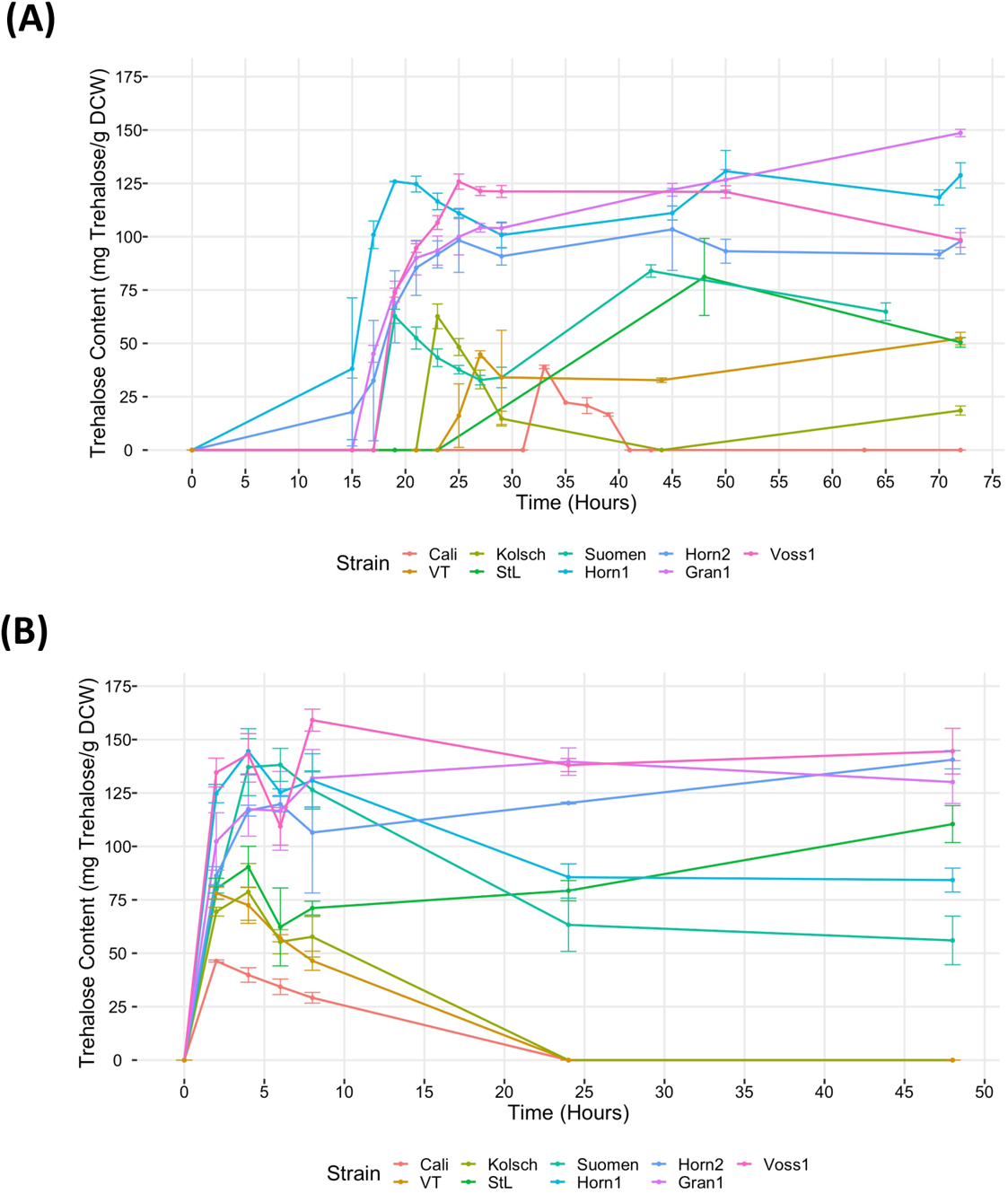
Analysis of trehalose accumulation during growth in YPD at (A) 30°C and (B) 37°C. Single colonies were inoculated into 100 mL of YPD and incubated at 30°C or 37°C with shaking at 170 rpm for 72 and 48 hours, respectively. Cell samples were collected at the indicated times and trehalose extractions were performed and concentrations determined via HPLC (see Methods). Data points represent the mean of biological replicates (n=3), and error bars represent the standard deviation.

Yeast cells are known to synthesize trehalose in post-diauxic (respiratory) growth followed by its hydrolysis into stationary phase (Lillie and Pringle, 1980; Parrou et al., 1999). The Kölsch and Cali controls displayed a rapid hydrolysis of its accumulated trehalose. The trehalose content in Kölsch and Cali peaked at 65 and 40 mg trehalose/g DCW and was completely consumed within the next 21 and 8 hours, respectively (Figure 6A). Also, subsequent prolonged growth of these control strains displayed minimal levels of trehalose being maintained within Cali, while Kölsch displayed a slow accumulation of trehalose as the cells progressed further into stationary phase. Suomen Hiiva and Vermont rapidly hydrolyzed approximately half of the synthesized trehalose, before switching to accumulating trehalose again (Figure 6A). By contrast, the kveik strains not only produced much more trehalose faster than the controls, but also consistently maintained high levels of trehalose throughout prolonged growth. While Hornindal1, Hornindal2 and Voss1 showed a reduction of its initial intracellular trehalose shortly after synthesis (~20%, ~5%, and ~8%, respectively), trehalose accumulation in the Hornindal strains recovered to its original levels during prolonged growth (Figure 6A). Voss1, on the other hand, showed a noticeable decline into stationary phase. Interestingly, Granvin1 showed no reduction in intracellular trehalose levels, but strikingly continued to accumulate trehalose throughout post-diauxic growth into stationary phase where it ultimately reached a maximum concentration of 148 mg trehalose/g DCW after 72 hours, equivalent to nearly 15% of the dry cell weight. As higher concentrations of intracellular trehalose support several mechanisms of cellular protection against environmental stresses it stands to reason that this enhanced trehalose accumulation in kveik strains could contribute to their survival and observed increased fermentation capabilities at higher temperatures.

When yeast strains were grown to late exponential phase at 30°C and transferred to 37°C for further growth, trehalose accumulated to high levels within the first 2 hours and peaked within 6 hours of heat treatment (Figure 6B). The Beer 1 control yeasts accumulated trehalose to lower levels, with Cali reaching ~46 mg trehalose/g DCW and Vermont, Kölsch and St. Lucifer almost double that amount (~78-90 mg trehalose/g DCW). Cali, Vermont and Kölsch degraded the accumulated trehalose to levels below detection within the next 20 hours of heat exposure. However, St. Lucifer, the more heat tolerant Beer 2 control strain, showed ~33% reduction in trehalose after 4-6 hours of heat exposure, followed by a steady increase in trehalose accumulation to ~110 mg/g DCW. By contrast, the kveik strains and Suomen Hiiva accumulate trehalose within 6 hours of the heat treatment in the ~120 mg/g DCW (Hornindal2 and Granvin1) to ~140 mg/g DCW range (Hornindal1, Voss1 and Suomen Hiiva) (Figure 6B). Furthermore, Voss1, Granvin1 and Hornindal2 maintain higher levels of trehalose (105-145 mg/g DCW), while Hornindal1 and Suomen Hiiva, on the other hand, degraded between 40-50% of its accumulated trehalose by 48 hours into the heat treatment. In combination, these data show high levels of trehalose accumulation by kveik and the heat tolerant St. Lucifer strains after 48 hours at 37°C, which could contribute to their tolerance to heat treatment.

Trehalose biosynthesis in *S. cerevisiae* is controlled by the Trehalose Synthase Complex (TSC), which is composed of: the enzymes Tps1, which catalyzes the formation of trehalose-6-phosphate (T6P), and Tps2, which dephosphorylates T6P into trehalose; and two regulatory subunits, Tps3 and Tsl1 (Thevelein and Hohmann, 1995; Ferreira et al., 1996; Francois and Parrou, 2001). During stress conditions such as nutrient starvation or exposure to heat, *TSL1* expression is induced and it is responsible for assembling the complex and activating Tps1, while unphosphorylated Tps3 activates Tps2 for the complex to start synthesizing trehalose (Francois and Parrou, 2001; Trevisol et al., 2014). All six kveik strains possess SNPs in genes known to be involved with trehalose biosynthesis (Supplementary Table S2). While *TPS1* and *TPS2* contain isolated SNPs unique to the kveik strains, SNPs were more abundant in the regulatory subunits. Specifically, *TPS3* have four unique SNPs of which two are present in all strains; 2500A>G (Ser834Gly) is homozygous in all six strains, while 238G>A is homozygous in 3/6 strains (Supplementary Table S2). In addition, *TSL1* has 11 unique SNPs of which three are present in three or more of the kveiks strains.

The accumulation of intracellular trehalose is a balance between its biosynthetic and hydrolytic reactions (Singer and Lindquist, 1998a; Eleutherio et al., 2015). The hydrolysis of trehalose in *S. cerevisiae* can be carried out by three enzymes: the acid trehalase Ath1 and the neutral trehalases Nth1 and Nth2. Ath1 has an optimal pH of 5.0 and is mainly localized to the vacuoles during normal growth and stress conditions but is transported to the cell membrane during stress recovery to hydrolyze extracellular trehalose and the resultant glucose molecules are transported into the cell via hexose transporters (Jules et al., 2004; Jules et al., 2008). Ath1 has low activity levels during exponential growth on fermentable sugars and high activity levels during the stationary phase, suggesting that it is likely regulated via catabolite repression by glucose (Jules et al., 2008; Magalhaes et al., 2018). By contrast, neutral trehalases possess an optimum activity at pH 7.0, are localized in the cytosol, and are primarily responsible for intracellular trehalose degradation (Nwaka et al., 1995; Elbein et al., 2003; Magalhaes et al., 2018). *NTH1* and *NTH2* have low expression during the exponential phase on glucose, increased expression towards the end of late exponential phase and peak in expression at the onset of stationary phase (Magalhaes et al., 2018). The consistently high levels of trehalose detected in the kveik strains suggests that its neutral trehalase activity may be dysfunctional, resulting in a decrease in the ability to effectively degrade intracellular trehalose and leading to high levels of accumulation. We consequently collected samples for trehalase activity assays 2 hours after the maximum trehalose spike in post-diauxic growth. The Beer 1 control strains possessed significantly higher neutral trehalase activities ranging from 58 µmol/min/mg for Kölsch to 120 µmol/min/mg for Cali (Figure 7). This coincides with the rapid hydrolysis of trehalose after its initial maximum accumulation (Figure 6A). Strikingly, the kveik strains supported a substantial reduction in neutral trehalase activities, ranging from 12–22 µmol/min/mg (Figure 7), which is 10-18% of the activity supported by Cali. Interestingly, Suomen Hiiva, the trehalose accumulating control, had 11% of the trehalase activity of Cali (Figure 7), but still hydrolyzed 52% of the synthesized trehalose, before it again accumulated trehalose into stationary phase (Figure 6A). While the neutral trehalase activities of the kveik strains were fairly similar to that of Suomen Hiiva, the kveik strains accumulated substantially more trehalose (1.5 – 2-fold or more) into stationary phase. These observations suggest a deficient trehalase activity along with increased biosynthesis resulting in rapid accumulation and the maintenance of high levels of intracellular trehalose that could support increased fermentation efficiency, stress tolerance and cell survival of kveik.

**Figure 7.**
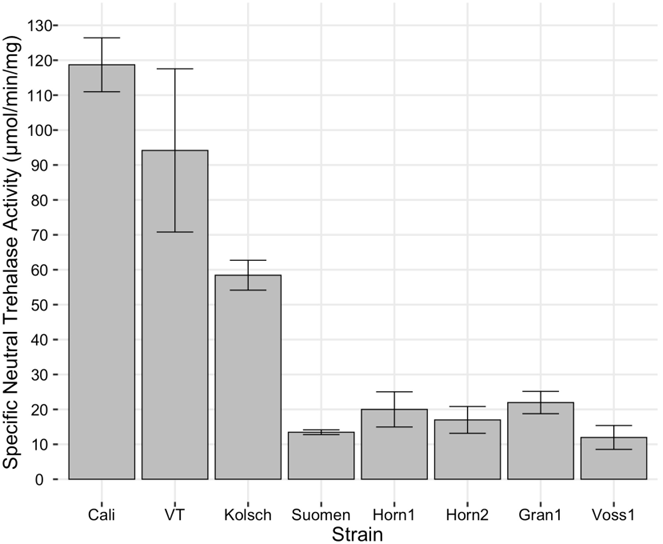
Analysis of specific neutral trehalase activity. Single colonies were inoculated into 100 mL of YPD and incubated at 30°C with shaking at 170 rpm until 2 hours post diauxic shift. Cell samples were collected and trehalase activity assays performed as described in Methods. Specific activity of trehalase is expressed as µmol of glucose liberated per min per mg protein. Data points represent the mean of biological replicates (n=3) and error bars represent the standard deviation.

We screened all three trehalases for potential mutations and identified several SNPs in all three genes (Supplementary Table S2). *NTH1* possesses several missense mutations near the catalytic site in Nth1 which could affect the folding and tertiary structure of the enzyme. In particular, 1213C>T (Leu405Phe) is homozygous in 4/6 kveik strains. Also, *NTH2* carries three unique homozygous SNPs (269G>C, Ser90Thr; 491C>A, Thr164Lys; and 1413T>A, Asn471Lys; Supplementary Table S2) in Granvin1, the strain that continuously accumulated trehalose in our assay (Figure 6). Lastly, *ATH1* carries seven unique SNPs, of which three are present in 3/6 or more kveik strains (Supplementary Table S2). These latter three mutations (3394A>G, Asn1132Asp; 3434A>G, Ser1145Asn; 3596A>G; Asn1199Ser) are either homozygous or present in high copy number in Granvin1. It is currently not known how these various SNPs and the coordinated functions of these genes impact trehalose accumulation and ultimately thermotolerance in the kveik strains.

## Discussion

Kveik are genetically distinct ale yeasts known for rapid fermentation at warmer temperatures while producing a diversity of interesting flavour compounds (Garshol, 2014; Preiss et al., 2018). Here we provide evidence of strain and temperature-specific variation in fermentation efficiencies and flavour compound production of six kveik strains at fermentation temperatures ranging from 12-42°C. These kveik strains also vary in their abilities to survive at higher temperatures. Strikingly, we identified significant increases in intracellular trehalose accumulation in kveik that could contribute its thermotolerance. Collectively, these findings provide further insight into the benefits and shortcomings of specific kveik strains at a range of different temperatures.

Ale fermentations are usually performed at 15-25°C, with temperatures around 20°C being the norm, while lagers are fermented colder (6-14°C) (Bamforth, 1998). Our fermentation analyses clearly identify a wide range of efficient fermentation temperatures for kveik strains with a preference for fermentations in the 30-37°C range (Figure 1). Strain variation is an important consideration as some strains, like Hornindal1, can efficiently attenuate at a broad temperature range (15-42°C), while others, like Laerdal2, prefer a narrower temperature range (30-37°C). The kveik strains in general attenuate wort faster than the control strains (other than Kölsch at 30°C) with FG ~ 1.01 already achieved after 48 hours at their fermentation temperature optima (Figure 1). This attribute is appealing as some strains, like Hornindal1 and Laerdal2, can be used for rapid beer production. Maltose constitutes ~60% of the fermentable wort sugar and its metabolism is repressed by glucose (Stewart and Russell, 1998; He et al., 2014). The rapid glucose consumption in kveik strains could relieve the repression on maltose metabolism, thereby allowing the faster initiation and ultimately complete consumption of maltose at a wider temperature range compared to the control strains (Supplementary Figures S2 and S3). The efficiency of maltose consumption is, however, strain and temperature dependent. Some strains, like Hornindal1, efficiently consume maltose at all temperatures tested while others, like Ebbegarden1, are less efficient at the temperature extremes tested (Supplementary Figures S2 and S3).

Maltotriose represents ~20% of the fermentable wort sugar and incomplete fermentation of maltotriose can be a major concern, especially in high- and very high gravity fermentations. Residual maltotriose causes process inefficiency (lower alcohol production from available sugar), a potential undesirable impact on flavour (Pachal and Stewart, 1979) and the added risk of undesirable post-packaging secondary fermentation (Krogerus et al., 2019). All the strains tested here showed incomplete maltotriose consumption. The Beer 1 controls were most efficient at maltotriose consumption at their respective fermentation temperature optima but struggled to consume this sugar outside these temperatures (Supplementary Figure S2). Kveik strains also failed to consume maltotriose completely; Hornindal1, Hornindal2 and Ebbegarden1 had <20% residual maltotriose at warmer temperatures, while the remaining strains generally left >25% of the maltotriose in the final beer. The amount of residual maltotriose in the final beer is both strain and temperature dependent (Supplementary Figures S2 and S3).

Higher alcohols such as propanol, butanol, isobutanol, and isoamyl alcohol, are often regarded as undesirable in beer and their production is induced by various fermentation stresses including higher temperatures (Saerens et al., 2008b). Preiss et al. (2018) previous reported isobutanol and isoamyl alcohol were produced below its sensory thresholds by kveik strains in 30°C fermentations. All the kveik strains tested here produced these higher alcohols at levels below the sensory threshold at all temperatures tested (Supplementary Table S1). This observation supports the use of these yeasts at a range of production temperatures without the risk of producing undesirable higher alcohols. Interestingly, we did notice 1-octanol, also found in citrus essential oils, was produced in a temperature-independent manner by most kveik strains and St. Lucifer at or slightly above its sensory threshold of 0.12 ppm at most temperatures, while only at cooler temperatures by Cali and Vermont (Supplementary Table S1). It is currently unclear how yeast would produce 1-octanol during fermentation. These findings provide brewers with a series of kveik yeasts with a broad range of brewing applications.

The kveik strains showed enhanced volatile fatty acid-related flavour metabolite production across most temperatures (Supplementary Table S1). MCFAs, like octanoic acid and decanoic acid, are known to be toxic to yeast cells and are potent fermentation inhibitors in a manner that is aggravated with increased acidity, ethanol and increasing temperature (Viegas and Sa-Correia, 1997). While both these MCFAs can negatively impact yeast cells, decanoic acid is more deleterious than octanoic acid (Viegas et al., 1989). Most of the kveik strains and St. Lucifer seems to have evolved mechanisms to circumvent MCFA toxicity as they maintain strong fermentation profiles at higher temperatures while producing higher levels of MCFA-associated flavour compounds. To this end, we detected very low levels of decanoic acid at the end of the fermentation if at all. Also, octanoic acid levels were noticeably higher at <30°C where it is less inhibitory and lower at >30°C where its toxicity increases. Yeast cells combat MCFA toxicity by controlling its synthesis, expelling the fatty acid from the cell and/or converting it to the much less inhibitory ethyl ester (Peddie, 1990; Piper et al., 1998; Cabral et al., 2001; Legras et al., 2010). To this end, whether the increased 1-octanol production in kveiks is a consequence of increased octanoic acid synthesis is not known. Some insight into these detoxification strategies can potentially be gleaned from the whole genome sequences previous reported for these strains (Gallone et al., 2016; Preiss et al., 2018). We identified several homozygous and heterozygous SNPs in key fatty acid metabolic genes (*AAC1*, *FAS1*, *FAS2* and *FAA1*) in several kveik strains (Supplementary Table S2). Whether these various SNPs contribute to an altered fatty acid metabolism in the kveik strains that would enable their growth and survival at elevated temperatures is currently not known.

Furthermore, two plasma membrane transporters known to be involved in various aspects of cellular detoxification, Tpo1 (polyamine transporter) and Pdr12 (ATP Binding Cassette transporter) (Piper et al., 1998; Hatzixanthis et al., 2003; Sa-Correia et al., 2009), have been identified as major components needed for the expulsion of octanoic and decanoic acids to combat their toxicity (Legras et al., 2010). In addition, *TPO1* expression increases as the fermentation proceeds, suggesting a role in fermentation-related stress responses (Marks et al., 2008). SNP analyses of *PDR12* and *TPO1* identified several heterozygous variants that are unique to either the kveik or control strains (Supplementary Table S2). Interestingly, two variants (*PDR1* 3622A>G, Ile208Val; and *PDR12* 1819A>G, Ile607Val) are homozygous in all the control strains, but heterozygous or unmutated in the kveiks strains. Also, *TPO1* carries a SNP (802G>C, Val268Leu) homozygous in some Beer 1 strains but absent from kveik strains. While the unique nature of these mutations are intriguing, the functional link between these variants and their impact on the abilities of Pdr12 or Tpo1 to export MCFAs from the cell is currently not known.

Lastly, ethyl ester production is known to increase as the fermentation temperature increases from 14-26°C (Saerens et al., 2008a). Esters are synthesized intracellularly and passively diffuse across the plasma membrane into the surrounding medium, with smaller molecules diffusing more rapidly (Nykänen and Nykänen, 1977). Longer chain ethyl esters, like ethyl decanoate, diffuse more slowly than the short acyl chain esters (i.e., ethyl hexanoate). The higher ethyl ester amounts at higher temperatures may be partially due to a more fluid and hence permeable plasma membrane. The kveik strains produced these fruity ethyl esters at above threshold concentrations at levels noticeably different than the controls and most abundant production was generally in the moderate temperature range (30-37°C) (Supplementary Figure S5). Laerdal2, Voss1 and Hornindal1 seemed to produce these ethyl esters more abundantly and at different temperatures than the other kveik strains. The efficiency of ethyl ester production is therefore strain and temperature dependent and allows for the production of a beer that has a fruity ester profile distinct from the Beer 1 controls. Ethyl ester synthesis was proposed as a mechanism of detoxifying the yeast cells of the inhibitory impacts of MCFAs (Peddie, 1990; Legras et al., 2010). For all yeasts tested we could only detect low amounts of decanoic acid. It is thought that the yeast cells prioritized converting decanoic acid to ethyl decanoate to rid the cell of the more toxic MCFA rapidly. Key genes in ethyl ester synthesis (*EEB1*, *EHT1* and *PLB2*) (Merkel et al., 1999; Saerens et al., 2006; Saerens et al., 2008a; Steyer et al., 2012) all carry several heterozygous mutations that are unique to either the kveik or the control strains (Supplementary Table S2), suggesting different capacities to produce ethyl esters. Whether either of these SNPs impact ethyl ester production warrants further investigation.

Our cell viability assay showed the Beer 1 control yeasts were more susceptible to heat treatment than St. Lucifer and the kveik strains (Figure 5). While the kveik strains generally have increased cell viabilities at higher temperatures, they are not equally thermotolerant as Hornindal1, Granvin1 and Voss1 are clearly more resistant to 40-42°C, while Hornindal2, Laerdal2 and Ebbegarden1 are more sensitive. As expected, the more heat tolerant strains also fermented more efficiently at higher temperatures than the more susceptible strains (Figure 1). Trehalose is known to protect yeast cells from a variety of environmental impacts, including cold, heat and ethanol stress (Coutinho et al., 1988; Eleutherio et al., 1993; Bandara et al., 2009; Eleutherio et al., 2015). Intracellular trehalose accumulation is controlled by the interplay between its biosynthesis, hydrolysis and export (Singer and Lindquist, 1998b; Eleutherio et al., 2015). Trehalose accumulation under standard growth conditions begins during the diauxic shift when glycogen reserves are being used and continues until stationary phase is reached when its hydrolysis proceeds into the rest of the stationary phase (Lillie and Pringle, 1980; Elbein et al., 2003; Eleutherio et al., 2015). However, in response to stress, including heat shock (37°C), genes implicated in trehalose production are induced resulting in the formation and activation of the TSC and concomitant trehalose biosynthesis, while its hydrolysis by trehalases are reduced to accumulate trehalose intracellularly where it is proposed to bind and stabilize proteins and membranes (Singer and Lindquist, 1998b; Francois and Parrou, 2001; Trevisol et al., 2014; Eleutherio et al., 2015). During stress recovery conditions, Protein Kinase A (PKA) phosphorylates Tps3 thereby inhibiting TSC activity. While all TSC genes carry SNPs, the regulatory genes *TPS3* and *TSL1* contained more variants. Specifically, the homozygous Ser834Gly and Val80Met variants in Tps3 and Ile101Val, Ile140Val, and Ile243Phe variants in the Tsl1 of Hornindal1 (Supplementary Table S2), the most stress tolerant kveik strain tested, suggests a deregulated TSC could result in the increased trehalose production observed in kveik (Figure 6). A higher internal concentration of trehalose protects cells against thermal stresses by supporting the plasma membrane and suppressing the aggregation of misfolded proteins (Eleutherio et al., 1993; De Virgilio et al., 1994; Hottiger et al., 1994). Although high concentrations of trehalose prevent proper refolding of proteins following a heat shock, constitutive production of significant quantities of trehalose could prime kveik for a variety of stresses and explain their ability to withstand high temperatures and adverse conditions better than their brewing yeast relatives.

Neutral and acidic trehalases hydrolyze trehalose. Neutral trehalases Nth1 and the weaker Nth2 have pH optima of 7.0 and function intracellularly to hydrolyze trehalose in the cytosol (Nwaka et al., 1995; Elbein et al., 2003; Magalhaes et al., 2018). By contrast, the acidic trehalase Ath1 has a pH optimum of 4.5, localizes to the vacuole and is transported to the plasma membrane to hydrolyze extracellular trehalose (Jules et al., 2004; Jules et al., 2008). Interestingly, the activities of the neutral trehalases and Ath1 differ temporally. The Nth1 is active during exponential growth in rich media, which decreases when glucose become limiting and is ultimately inactive in starvation (Thevelein, 1984; App and Holzer, 1989; San Miguel and Arguelles, 1994; Thevelein and Hohmann, 1995), while Ath1 is inactive in exponential growth, but becomes active along with Nth2 in stationary phase when the cells starve (Jules et al., 2008; Garre et al., 2009). The kveik strains accumulate substantially more intracellular trehalose in rich media than commercial Beer 1 ale strains (Figure 6). The significantly reduced neutral trehalase activities in post-diauxic growth of the kveik strains could be at least partially responsible for the increased intracellular accumulation of trehalose (Figure 6) and hence contributes to stress tolerance. While the molecular basis for the decreased neutral trehalase activity in kveiks remain unresolved, it is known that trehalases are controlled at both the gene transcription and enzyme activity levels. For example, the transcription of the *NTH1* is low in glucose (exponential growth) and it increases when glucose becomes limiting in the diauxic shift and into stationary phase (Zähringer et al., 1998; Parrou et al., 1999). In addition, *NTH1* transcription is also induced under different stress conditions (e.g., heat and osmotic stress) via STREs in its promoter by Msn2/Msn4 in a PKA-dependent manner (Zähringer et al., 1997; Zahringer et al., 2000). By contrast, Nth1 is active during exponential growth in abundant glucose, it decreases in the diauxic shift and is low in stationary phase cells (Thevelein, 1984; App and Holzer, 1989; San Miguel and Arguelles, 1994; Thevelein and Hohmann, 1995). PKA directly phosphorylates and activates Nth1 in exponential growth. The activity of Nth1 therefore decreases as glucose becomes limiting as PKA becomes inactive, leading to an increase in trehalose synthesis and accumulation (Schepers et al., 2012; Veisova et al., 2012; Eleutherio et al., 2015). It is currently unclear if the reduction in neutral trehalase activity in kveik is due to a lack of gene expression, reduced protein content, lack of phosphorylation/activation, or inefficient hydrolysis of trehalose due to the SNPs we identified in *NTH1* and *NTH2* (Supplementary Table S2).

In addition, the three kveik strains with the highest temperature tolerances, Hornindal1, Voss1 and Granvin1 (Figure 4), also contain a chromosomal aneuploidy of 4n+1 of chromosome IX (Preiss et al., 2018)(Supplementary Table S3). *BCY1*, the gene encoding the regulatory subunit for cAMP dependent PKA, is located on this chromosome. In the absence of cAMP, Bcy1 inhibits PKA by forming an inactive complex with the PKA catalytic subunits (Hixson and Krebs, 1980; Toda et al., 1987). When cAMP is present, it binds Bcy1 and causes its dissociation from the complex with PKA thereby releasing the catalytic subunits. Overexpression of *BCY1* is linked to reduced PKA activity which leads to a decrease in Nth1 activity and results in greater trehalose accumulation and increased thermotolerance (Portela et al., 2003). Increased expression of *BCY1* in these three kveik strains could contribute to the increased intracellular accumulation of trehalose and reduction in neutral trehalase activity.

Agt1, the main maltotriose transporter in beer yeast, is a broad substrate range α-glucoside transporter with high affinity for trehalose and sucrose (Km ~8 mM), and low affinity for maltose and maltotriose (K_m_ ~ 35 mM) (Han et al., 1995; Stambuk and de Araujo, 2001; Alves et al., 2007; Alves et al., 2008). Agt1 is also important to combat environmental stress (da Costa Morato Nery et al., 2008; Ratnakumar et al., 2011). This transporter is therefore pivotal in both maltotriose and trehalose metabolisms in beer yeasts. The *AGT1* genes of lager and ale yeasts have been sequenced and shown to be similar, but not identical to *MAL11* (Vidgren et al., 2005). Upon closer scrutiny of the *AGT1* genes isolated from lager and ale yeasts, the authors identified SNPs that are either unique to or shared by these brewing yeasts. Strikingly, a frame shift insertion mutation at position 1183 resulted in a premature STOP codon at Agt1 Glu395 present in the lager yeasts, but absent from the ale yeasts tested (Vidgren et al., 2005). Gallone et al. (2016) later reported this mutation as homozygous in many Beer 2 yeasts including St. Lucifer, the thermotolerant and maltotriose-fermentation deficient Belgian ale yeast. In addition, a 1643T>C SNP (Val548Ala) was found in ale yeast *AGT1*, specifically in transmembrane domain VIII (Vidgren et al., 2005) that was later shown to impact maltose transport at 20°C (Smit et al., 2008; Vidgren et al., 2014). Here we identified two additional heterozygous frameshift mutations and two gained STOP codon mutations in kveik (Supplementary Table S2), suggesting heterozygous loss-of-function of *AGT1* in 4/6 kveik strains. With premature STOP codons previously identified in lager yeasts, the presence of four such loss-of-function mutations in these traditional Norwegian farmhouse ale yeasts could potentially represent a mechanism of evolutionary adaptation to challenging environmental pressures, such as repeated cycles of fermentation at elevated temperatures followed by desiccation experienced during its traditional usage (Garshol, 2014). Furthermore, there are several other SNPs in the kveik *AGT1* genes that were previously identified in lager yeast or that was not reported by Vidgren et al. (2005) (Supplementary Table S2). The impact(s) of these missense mutations on Agt1 function is currently unknown. While these kveik strains phylogenetically form a unique subpopulation of Beer 1 ale yeasts, phased haplotypes previously revealed an admixed ancestry with contributions from outliers in Beer 1 as one haplotype and the other with contributions from diverse origins (Preiss et al., 2018). The origins and larger functional implications for these mutations in *AGT1* in kveik are therefore unknown.

As an high-affinity trehalose H^+^-symporter, the role of Agt1 in combatting environmental stress (da Costa Morato Nery et al., 2008; Ratnakumar et al., 2011) is proposed to export intracellularly accumulated trehalose to the periplasmic space where it binds the polar heads of phospholipids in the outer leaflet of the plasma membrane to protect it from environmental stresses (Eleutherio et al., 1995; Magalhaes et al., 2018). We have shown that the strains with the greatest intracellular trehalose content (Hornindal1, Granvin1, and Voss1) maintained significantly higher rates of viability after growth at 42°C (between 80-90% viability) compared to the Beer 1 strains that accumulate less trehalose. Kveik yeasts have also accumulated a variety of SNPs in their *AGT1* genes that have not yet been functionally characterized. While it is currently unknown if any of these mutations impact thermotolerance, it is noteworthy that the strains accumulating loss-of-function *AGT1* SNPs (Ebbegarden1, Laerdal2 and Hornindal2) were more sensitive to heat treatment (Supplementary Table S2; Figure 5). Agt1 transports α-glucosides from the environment into the cytosol (Stambuk et al., 1998; Stambuk and de Araujo, 2001; Alves et al., 2008) and also exports trehalose from the cytosol to the external environment (periplasm) (Eleutherio et al., 1995; Eleutherio et al., 2015; Magalhaes et al., 2018). Magalhaes et al. (2018) proposed Agt1 to be a reversible transporter, much like Mal61 for maltose, Fps1 for glycerol and SLC1 (an active transporter) for glutamate (Luyten et al., 1995; van der Rest et al., 1995; Grewer et al., 2008). In such a scenario, kveik strains would be challenged to export trehalose with high affinity to protect against thermal and ethanol stress for survival, while also importing maltotriose from the environment with low affinity to consume the last available fermentable sugar in the wort. With Agt1 function potentially compromised in kveik strains (and St. Lucifer) through heterozygous loss-of-function mutations, this might explain why these yeasts have high residual maltotriose in the beer at the end of the fermentation.

Overall, the kveik SNPs we report here provide some insight into the genetic adaptations in genes relevant to various different aspects of fermentation. To fully understand the impacts of these genes on the respective fermentation phenotype, gene expression analyses and functional assays of the relevant allele would be needed. Such experiments would be the focus of studies to link these kveik genotypes to a molecular understanding of the respective phenotypes.

Since their introduction to the scientific literature in 2018, traditional Norwegian kveik yeasts have become extremely popular in the brewing industry, especially in applications with craft breweries and home brewers. The present work extensively demonstrates the variation in kveik strains at a wide range of temperatures and shows that not all kveik strains are equal in terms of fermentation performance, stress tolerance, and flavour metabolite production. The data presented can help brewers to select optimal temperature and yeast strains to optimize their beer flavour and performance outcomes. Of relevance to brewers, kveiks are phylogenetically a part of Beer 1 (Preiss et al., 2018) with rapid glucose and maltose metabolism. However, kveiks share several phenotypic characteristics with Beer 2 yeasts, such as St. Lucifer, including temperature tolerance and flavour metabolite intensity (Calahan et al., 2011; Tapia et al., 2015).

This work also presents several opportunities for future development. Here, we showed that kveik is a suitable option for fermentation at high temperatures with potential applications not only for beer production but also in other high-temperature fermentation industries such as distillation or biofuels. The surprising presence of 1-octanol among the kveik yeasts and St. Lucifer strain may support the use of these yeasts as platform strains for the development of biofuels. The ability to fine-tune fatty acid ester aroma production in food fermentation yeasts remains a mystery, and the polymorphism of MCFA metabolic genes in kveik may present an opportunity to better understand fatty acid ester production. In this paper, we demonstrate that kveik show enhanced trehalose accumulation, which confers heat tolerance. This is likely linked to polymorphisms in *AGT1*, which may suggest a tradeoff between trehalose accumulation and maltotriose consumption. Trehalose accumulation is also known to improve desiccation tolerance, which is a valuable property for active dry yeast (ADY) production. Taken together our results present a clearer picture of the future opportunities presented by Norwegian kveik yeast as well as offer further insight into their applications in brewing.

## Author Contributions

BF, CT, RP, KK, ML, EO, and GV conducted the experiments and data analyses described in this study. KK, ML, and EO performed bioinformatic analysis of the whole genome sequence data. BF, CT, RP and GV designed the experiments. BF, CT, EO, RP, KK and GV prepared the manuscript. All authors read and approved the final manuscript.

## Funding

This research was funded by Natural Sciences and Engineering Research Council of Canada Discovery (NSERC #264792-400922), OMAFRA-University of Guelph Gryphons LAAIR Product Development (UG-GLPD-2020-100357), and Ontario Regional Priorities Partnership program (ON-RP3 06) grants.

## Acknowledgements

We thank Marvin Dyck and Wellington Brewery for donating the wort used in this research.

## Conflict of interest

Richard Preiss was employed by Escarpment Laboratories Inc. Kristoffer Krogerus was employed by VTT Technical Research Centre Ltd. The authors declare that the research was conducted in the absence of any commercial or financial relationships that could be construed as a potential conflict of interest.

## Notes

### Competing Interest Statement

The authors have declared no competing interest.

