## Supplementary Figure S1 for "Kveik brewing yeasts demonstrate wide flexibility in beer fermentation temperature and flavour metabolite production and exhibit enhanced trehalose accumulation"

**(A)**

| Stain name | Source | References |
|---|---|---|
| Cali Ale | Escarpment Laboratories | Preiss et al., 2018 |
| Vermont Ale | Escarpment Laboratories | Preiss et al., 2018 |
| Kölsch | Escarpment Laboratories | Gallone et al., 2016 |
| St. Lucifer | Escarpment Laboratories | Preiss et al., 2018 |
| Hornindal 1 | Escarpment Laboratories | Preiss et al., 2018 |
| Hornindal 2 | Escarpment Laboratories | Preiss et al., 2018 |
| Voss 1 | Escarpment Laboratories | Preiss et al., 2018 |
| Granvin 1 | Escarpment Laboratories | Preiss et al., 2018 |
| Stordal Ebbegraden 1 | Escarpment Laboratories | Preiss et al., 2018 |
| Laerdal 2 | Escarpment Laboratories | Preiss et al., 2018 |
| Suomen Hiiva | Suomen Hiiva | Katina et al., 2007 |

**(B)**

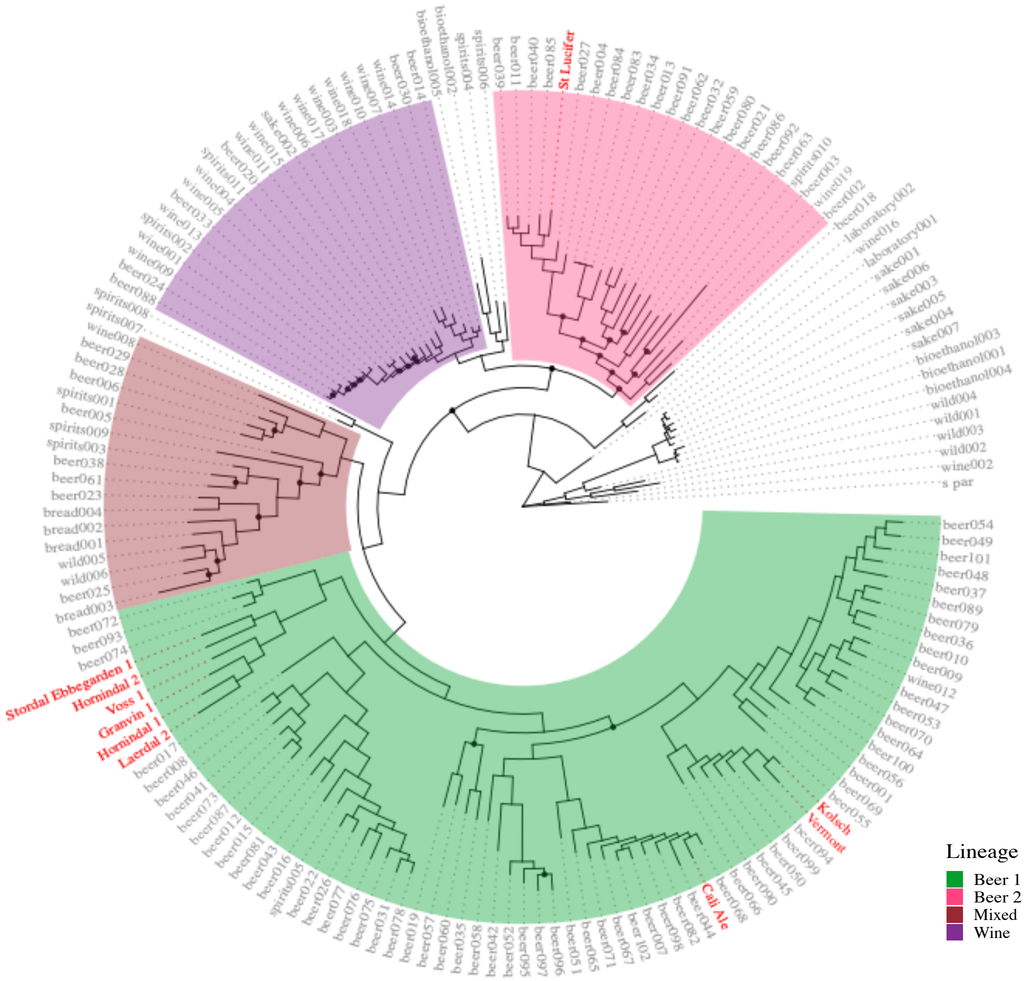


**Supplementary Figure S1. Yeast strains used in this research.** (A) Sources of the yeast strains used. (B) Phylogeny of the strains used compared to the *S. cerevisiae* strains sequenced in Gallone et al. (2016). The four 'main' lineages from Gallone et al. (2016) are colour-coded and the strains used in this study are shown in red.
