## Supplementary Figure S2 for "Kveik brewing yeasts demonstrate wide flexibility in beer fermentation temperature and flavour metabolite production and exhibit enhanced trehalose accumulation"

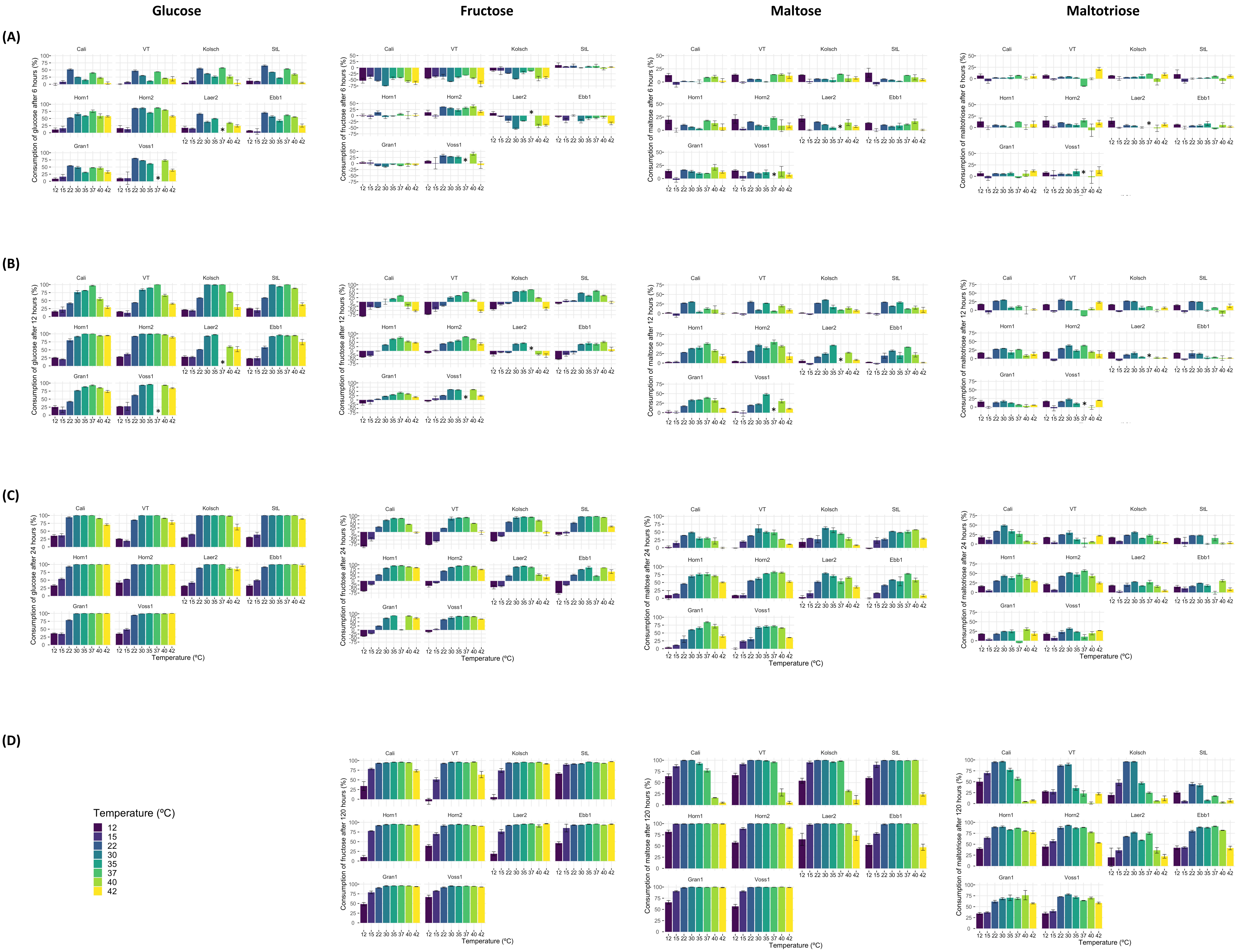


**Supplementary Figure S2.** Wort sugar consumption during the fermentation in Figure 1. Samples were collected at 6h (A), 12h (B), 24h (C), and 120h (D) of fermentation. The concentrations of glucose, fructose, maltose, and maltotriose were determined by HPLC as described in Methods. Data points represent the mean of biological replicates (n=3) and error bars represent the SD. * sugar concentrations not determined (sample processing inconsistencies).
