## Supplementary Figure S3 for "Kveik brewing yeasts demonstrate wide flexibility in beer fermentation temperature and flavour metabolite production and exhibit enhanced trehalose accumulation"

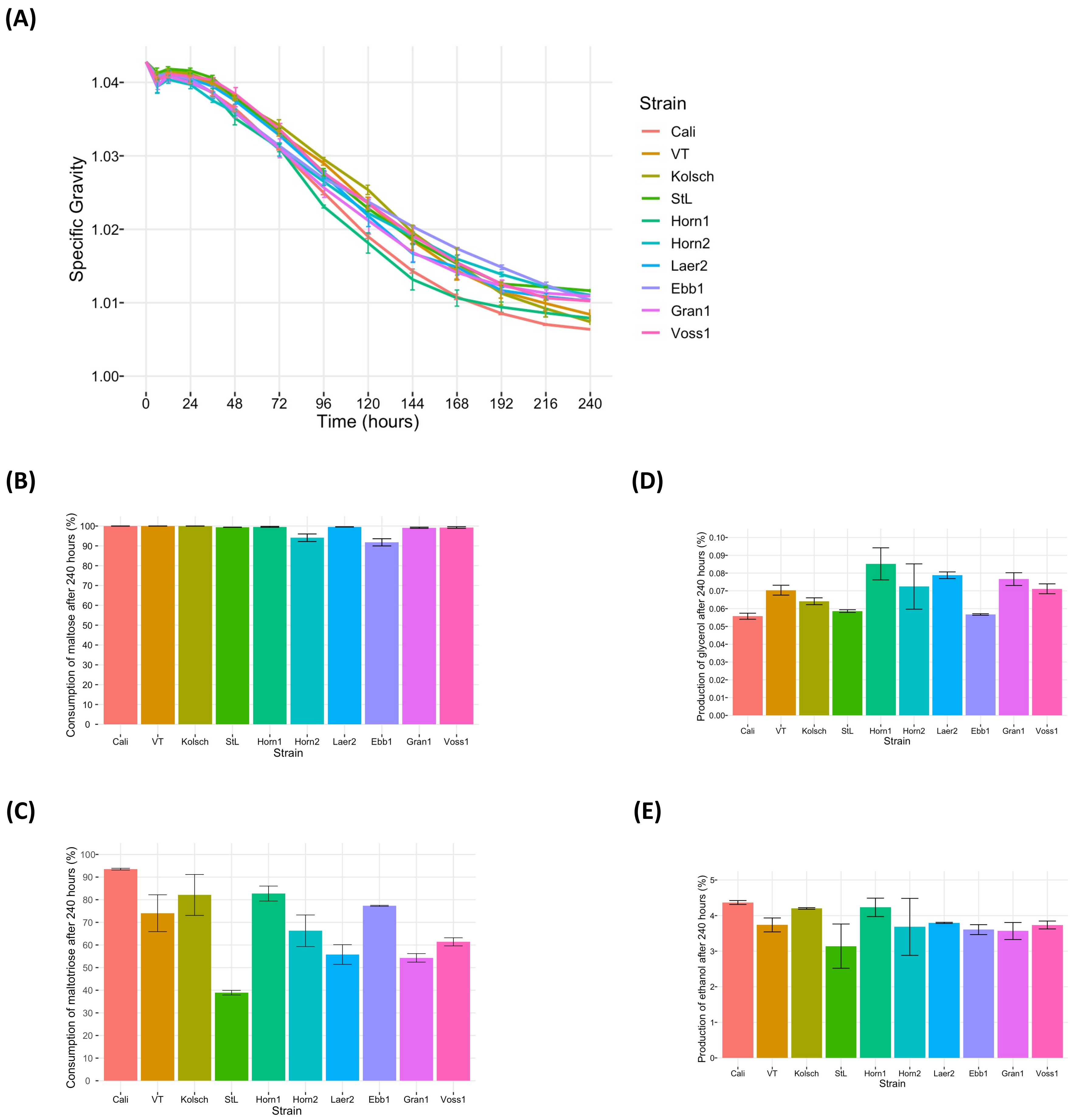


**Supplementary Figure S4.** Fermentation profiles (A), wort sugar consumption (B, C), and metabolite production (D,E) of four commercial *Saccharomyces cerevisiae* beer strains and six Norwegian kveik isolates during a prolonged cold fermentation at 12°C for 10 days. Strains were pre-cultured and wort inoculated as described in the Methods. Fermentation profiles were obtained via analyzing the change in specific gravity throughout fermentation using a DMA35v4 portable densitometer (Anton-Paar). Metabolic data was generated through high performance liquid chromatography (HPLC) as described in Methods. Data points represent the mean of biological replicates (n=3) and error bars represent the SD.
