## Supplementary Figure S4 for "Kveik brewing yeasts demonstrate wide flexibility in beer fermentation temperature and flavour metabolite production and exhibit enhanced trehalose accumulation"

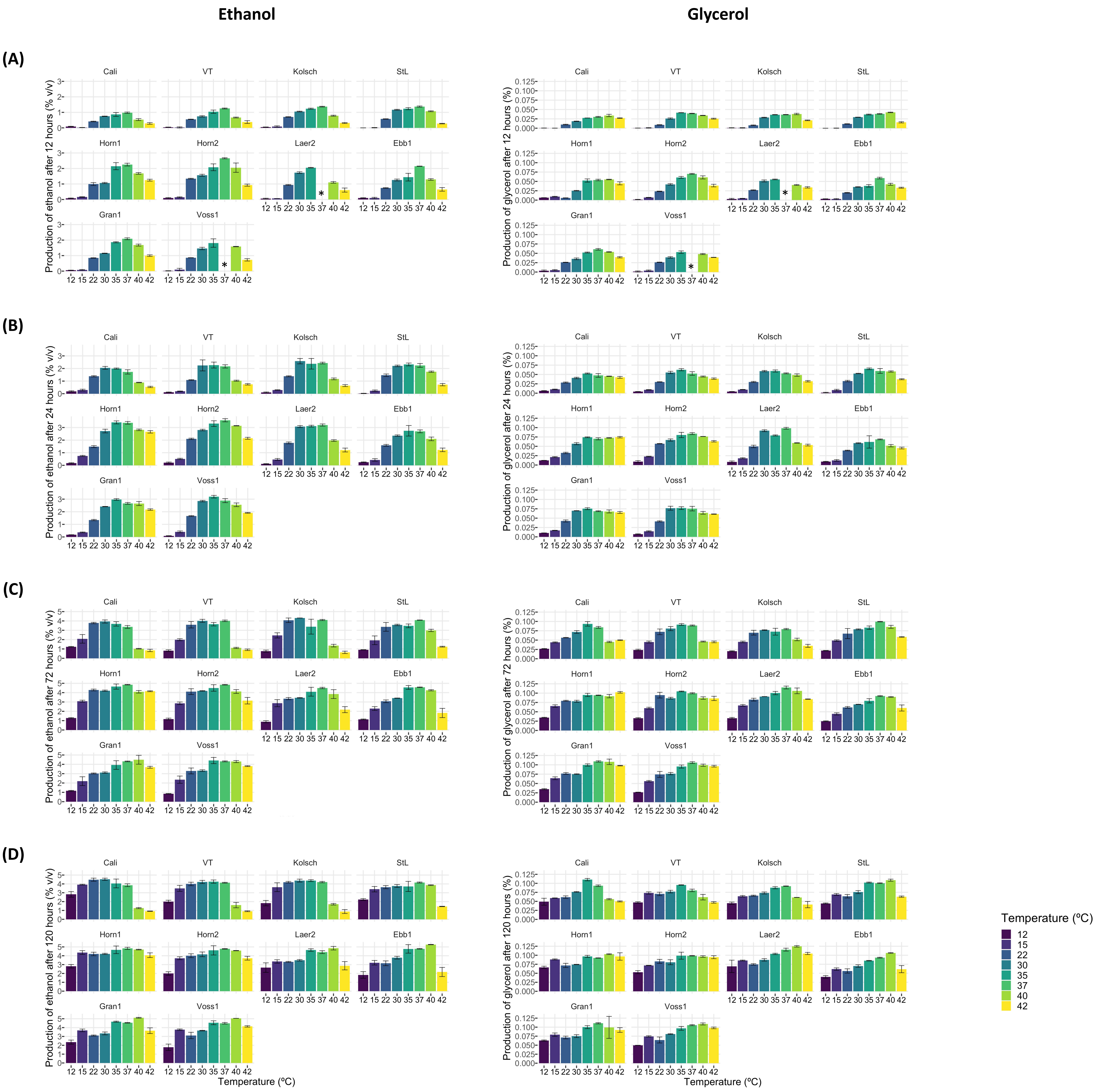


**Supplementary Figure S4.** Ethanol and glycerol production of six Norwegian kveik strains and four commercial *Saccharomyces cerevisiae* beer strains during the fermentation in Figure 1. Samples were collected for HPLC analyses at the same timepoints of SG measurements in Figure 1. The concentrations of ethanol and glycerol were determined at 12 h (A), 24 h (B), 72 h (C), and 120 h (D) by HPLC as described in Methods. Data points represent the mean of biological replicates (n=3) and error bars represent the SD. *metabolite concentrations not determined (sample processing inconsistencies).
