## Supplementary Figure S5 for "Kveik brewing yeasts demonstrate wide flexibility in beer fermentation temperature and flavour metabolite production and exhibit enhanced trehalose accumulation"

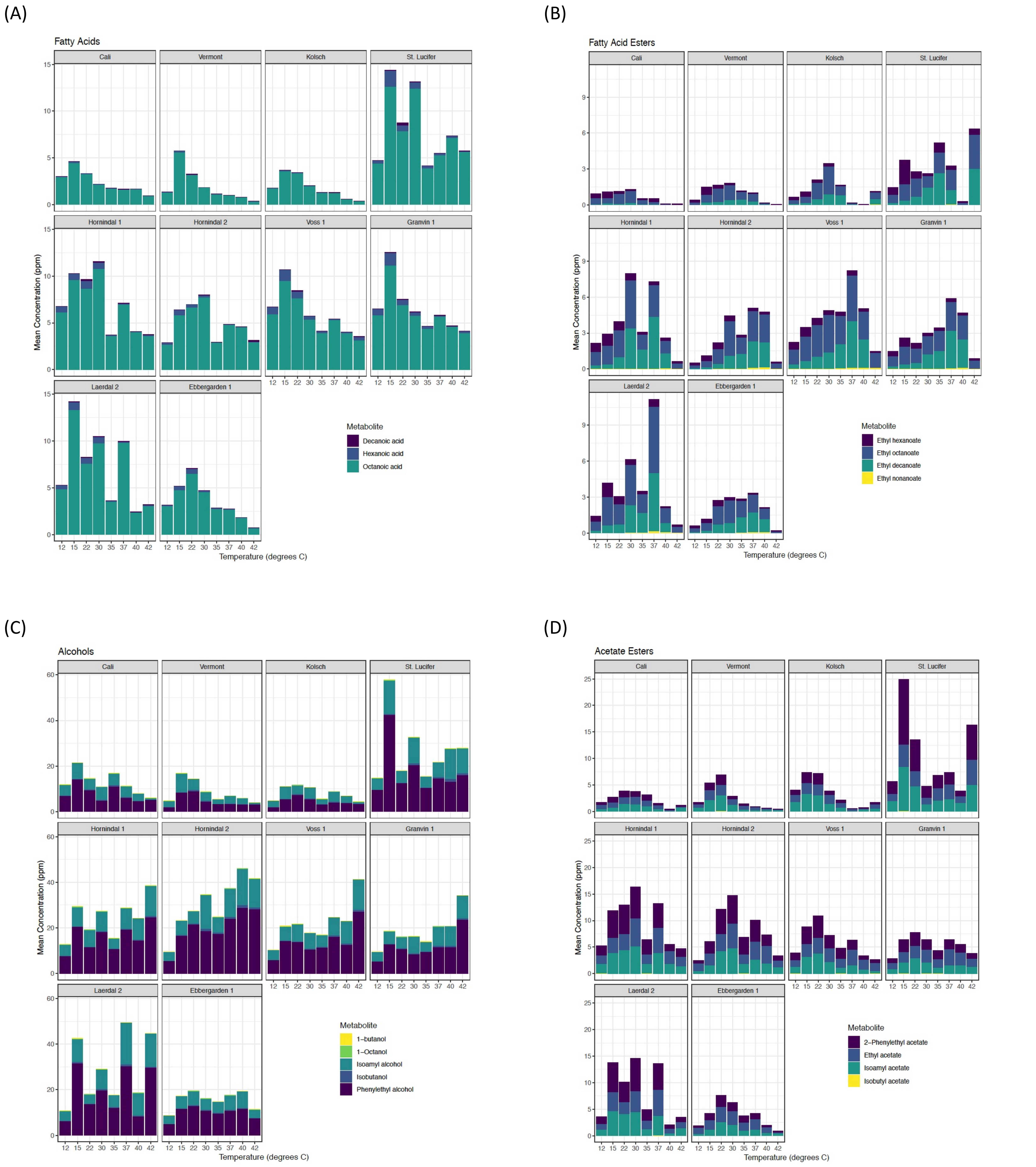


**Supplementary Figure S6.** Volatile flavour metabolite production of numerous fatty acids (A), fatty acid esters (B), higher alcohols (C), and acetate esters (D) by four control ale strains (Cali, Vermont, Kolsch, and St. Lucifer) and six kveik strains (Hornindal 1, Hornindal 2, Voss 1, Granvin 1, Laerdal 2, and Ebbegarden 1). Compounds were measured using HS-SPME-GC-MS. Final concentrations were determined through comparison with an internal standard of a known concentration. Data points represent the mean of biological replicates (n=3).
